# Calibration-free estimation of field dependent aberrations for single molecule localization microscopy across large fields of view

**DOI:** 10.1101/2024.12.11.627909

**Authors:** Isabel Droste, Erik Schuitema, Sajjad Khan, Stijn Heldens, Ben van Werkhoven, Keith A. Lidke, Sjoerd Stallinga, Bernd Rieger

## Abstract

Image quality in single molecule localization microscopy (SMLM) depends largely on the accuracy and precision of the localizations. While under ideal imaging conditions the theoretically obtainable precision and accuracy are achieved, in practice this changes if (field dependent) aberrations are present. Currently there is no simple way to measure and incorporate these aberrations into the Point Spread Function (PSF) fitting, therefore the aberrations are often taken constant or neglected all together. Here we introduce a model-based approach to estimate the field-dependent aberration directly from single molecule data without a calibration step. This is made possible by using nodal aberration theory to incorporate the field-dependency of aberrations into our fully vectorial PSF model. This results in a limited set of aberration fit parameters that can be extracted from the raw frames without a bead calibration measurement, also in retrospect. The software implementation is computationally efficient, enabling fitting of a full 2D or 3D dataset within a few minutes. We demonstrate our method on 2D and 3D localization data of microtubuli and nuclear pore complexes over fields of view (FOV) of up to 180 μm and compare it with spline-based fitting and a deep learning based approach.

## Main text

Single-molecule localization microscopy (SMLM) [Betzig2006, Rust2006, Sharonov2006, Heilemann2008] enables routine imaging at the nano-scale. The need for high throughput and much data implies that a large Field Of View (FOV) must be imaged, which is enabled by modern sCMOS sensors that have up to 10 Mpixels and advanced illumination schemes [Douglass2016, Deschamps2016, Diekmann2017, Archetti2019, Mau2021, Li2022, Ma2024, Nelson2024]. A key problem that arises is that aberrations depend on the position in the FOV. This is problematic for both the precision and accuracy of in-plane (*xy*) localization, but even more so for 3D (*xyz*) localization. Up to now aberrations, let alone field dependent aberrations, are rarely considered in SMLM as it is time consuming and cumbersome to i) measure them and ii) include them into the fitting. For that reason, the default method for estimating the positions of the emitters is to fit a simplified Point Spread Function (PSF) model to the recorded data. Typically, this simplified model is a Gaussian [Smith2010], which reduces the computational load to estimate the positions of millions of emitters to reconstruct a super-resolution image. Known drawbacks, however, are position biases in case of asymmetric aberrations and of emitters with partially fixed orientation [Stallinga2010], and an underestimation of the photon count [Thorsen2018]. An actual data driven PSF model is the spline model [Li2019], which requires calibration of the PSF by beads, potentially at various field and depth positions.

Advantageously, a physically correct fully vectorial PSF model could be used, so that high NA and polarization effects are properly considered. Moreover, (field dependent) aberrations of the microscope can then automatically be incorporated, and emission dipole orientation as well, if needed. Up to now such a full vectorial model is not used in practice for SMLM due to the computational load [Hullemann2021].

The current practical standard to measure field dependent aberrations is a calibration in the form of an axial scan of many fluorescent beads distributed over the whole FOV [Li2019, Hulleman2021, Fu2023]. An alternative is provided by the localization data itself, as millions of localization events are in fact millions of measurements of the microscope’s PSF across the FOV. The localization data itself thus provides a wealth of information on the optical system, and the challenge is to unlock that information. The key obstacle is that each individual localization event does not allow for fitting of aberration parameters next to the position, photon count and background, because the data is much too noisy for a robust estimation of many parameters. Previously, Xu et al. applied pupil phase retrieval to an averaged set of 3D localization data, but only so using an approximate scalar PSF model and for global aberrations independent of the position in the FOV [Xu2020]. Recently, Liu et al. proposed a framework for inverse PSF modelling, enabled by automatic differentiation, and showed its applicability to vectorial PSF fitting of 3D localization datasets with field dependent aberrations [Liu2024]. Here, we present an alternative method to estimate the field dependent aberrations from 2D or 3D single molecule images alone, without any calibration measurement. We make this possible by fitting the localization data with a global optical aberration model, with a limited number of fit parameters, instead of fitting the full aberration content for each measured PSF separately. This global aberration model is derived from the so-called Nodal Aberration Theory (NAT) [Shack1980], which provides explicit relations of the field dependency of the appearing aberration coefficients via low-order polynomials of the field coordinates. This eliminates the need for ad-hoc smoothing of field dependent aberrations as done in the inverse PSF modeling approach and enables application to 2D localization data.

Aberrations can be modelled by expressing the phase aberration function in the pupil plane in terms of Zernike modes. Using NAT, the Zernike coefficients *A*_*nm*_ can be expressed as low-order polynomials of the field coordinates (*x, y*) (see Supplementary Note for theory):

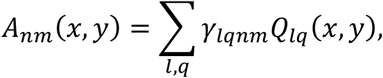

where *γ*_*lqnm*_ are the NAT coefficients, and *Q*_*lq*_(*x, y*) are products of (Legendre) polynomials of order *l* and *q*. We adapted NAT to the square shape of the FOV by using Legendre polynomials to describe the field dependence. For describing the field dependency of the first order aberrations, that is, defocus, and primary astigmatism, coma, and spherical aberration, it turns out that only 13 NAT coefficients are needed. To include second order aberrations (trefoil and secondary astigmatism, coma, and spherical aberration), an additional 43 NAT coefficients are needed. This number of free parameters, though large, is still orders of magnitude less than the number of free parameters in data driven approaches [Li2018, Bates2022].

Figure 1 schematically depicts our method. The raw blinking time series data is segmented into regions of interests (ROI) containing fluorescent emitters as usual for SMLM. From these millions of ROIs, a very small subset of *M*_*s*_ ∼ 10^3^ ROIs across the FOV is randomly selected to be used as input for the field dependent aberration estimation. The estimation process consists out of an alternation of local and global updates. During a global update, the NAT coefficients are updated while keeping the locations, photon count and local background of the emitters fixed, while during a local update, the locations are updated with fixed NAT coefficients. Subsequently, all ROIs are fitted with a vector PSF model (“Vectorfit”, https://gitlab.tudelft.nl/imphys/ci/vectorfit) using the found field dependent aberrations as input. We have implemented the vector PSF fitting on GPU for speedup and have devised several algorithmic improvements for additional steps in efficiency. First, we utilize the phasor method to provide a fast, initial estimate of the lateral position that is robust against high background [Martens2018]. Second, we compute the initial estimate of photon count *N*_*ph*_ and background count *bg* with linear regression given the data and the model. Third, pre-computing the Optical Transfer Function (OTF) reduces the number of needed Fourier Transformations per iteration by a factor of 6. These different algorithmic innovations are outlined in the Methods section.

**Fig. 1.**
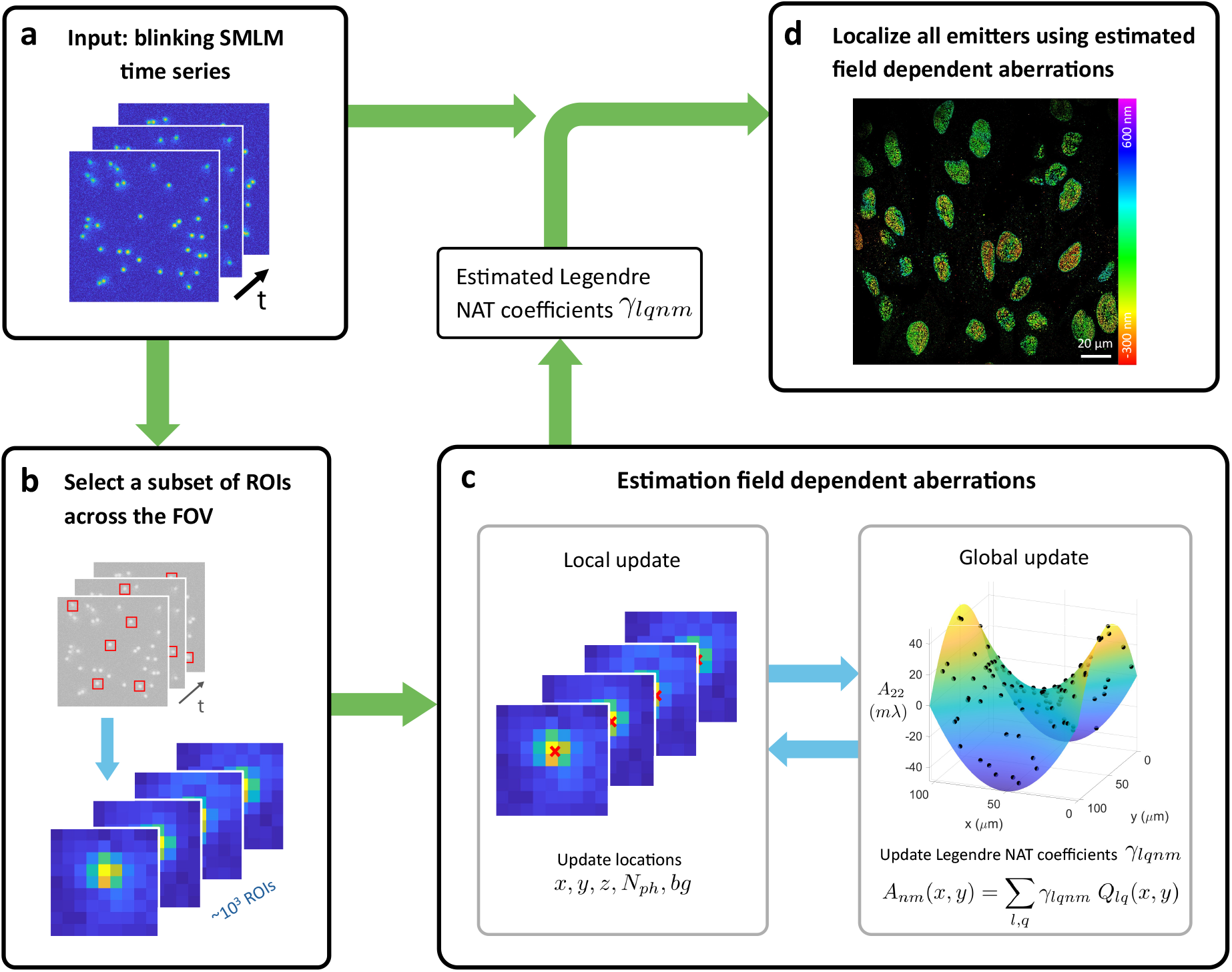
Schematic of fitting field dependent aberrations from single molecule data. **a**, The raw single molecule blinking frames are used as input. **b**, After segmentation into ROIs, a subset of ∼10^3^ ROIs across the FOV is selected for field dependent aberration estimation. **c**, The estimation of field dependent aberrations uses maximum likelihood estimation and consists of two optimization loops: a local update that updates the locations, photon count and background of each ROI while keeping the NAT coefficients constant, and a global update that updates the Legendre NAT coefficients while keeping the locations, photon count and background constant. By selecting different ROIs in **b** multiple times and repeating the estimation (**c)** an estimation precision for the aberrations can be calculated. **d**, Finally, in all ROIs emitters are localized using the vectorial PSF model (Vectorfit) with the estimated NAT coefficients as input.

We have tested our method on SMLM data of microtubuli acquired over a large 97 × 97 µm FOV (see Methods for experimental details). The same sample was imaged with and without astigmatism to obtain 2D and 3D localization data, respectively (see Figure 2). The aberrations found from the single molecule data via our method give very similar aberration maps across the FOV (orange in Fig. 2a,e) as the aberration maps obtained by interpolating between aberrations retrieved from *z*-stack bead calibration data (light blue in Fig. 2a,e), with the largest difference in the slope of the coma aberration maps. We performed a chi-square goodness of fit test [Siemons2018] to check if the PSF model with single molecule derived aberrations gives rise to a better fit than the PSF model with bead calibration derived aberrations (Supplementary Figure 1). For both the 2D and 3D data we find on average about 1% smaller chi-square values for the single molecule derived aberrations, up to about 5% and 3% smaller at the edges of the FOV for the 2D and 3D datasets, respectively, indicating a slightly better fit of the single molecule derived aberrations. The overall chi square values are about 20% (2D) and 16% (3D) higher than the theoretical values based on shot noise statistics [Siemons2018] (Supplementary Figure 3), indicating a residual model mismatch, possibly due to higher order aberrations and amplitude aberrations. A further validation of our approach is found by a comparison of the modelled PSF to the measured single molecule spots in 4×4 patches across the FOV, as a function of the axial position (Supplementary Video 1). This comparison shows that the modelled PSF matches very well with the measured data.

**Fig. 2.**
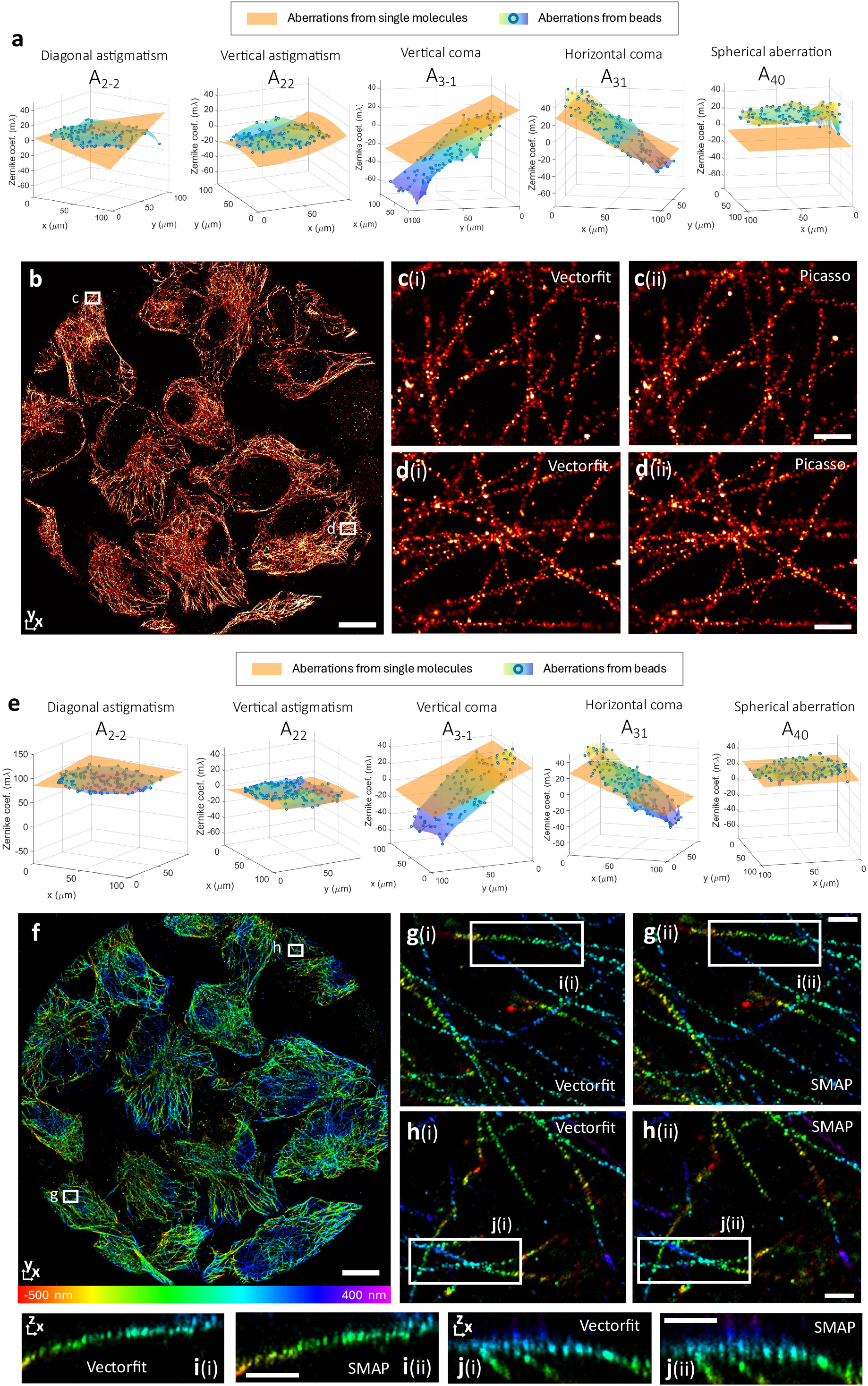
Aberration estimation from 2D and 3D experimental data of microtubili. A sample was imaging in 3D with and in 2D without astigmatism. **a**, Estimated Zernike aberration surfaces from single molecules of the 2D data, compared to interpolated aberrations from bead z-stack calibration. **b**, Full FOV image reconstructed using Vectorfit and fitted aberrations from single molecules. **c**, and **d**, insets of **b** where (i) shows vector fitting and (ii) Gaussian fitting by Picasso [Picassco] **e**, Estimated Zernike aberration surfaces from single molecules of 3D data, compared to interpolated aberrations from bead z-stack calibration. **f**, Full 3D FOV image reconstructed using Vectorfit and fitted aberrations from single molecules. The 3D image is shifted 14 µm to the upper right compared to the 2D image. The false color indicates the axial distance from the nominal focus. In the 3D image, the microtubules that overlay the nucleus are visible in blue, while in the 2D image, these regions appear black, as the emitters are too much out of focus. **g**,**h** insets of **f** where (i) shows the Vectorfit result and (ii) with SMAP [SMAP]. **i**,**j**, *x*z-cross sections of the regions indicated in **g**,**h**. Scale bars: 10 μm (**b**,**f**), 500 nm (**c**,**d**,**g**,**h**,**i**,**j**)

We also found a difference in the estimated bead aberrations between consecutive days of measurement with different bead samples. The beads measured on the second day show a uniform shift in coma coefficient *A*_3−1_ (Supplementary Figure 2). By estimating the *z*-locations of the beads, we found that the tilt of the sample was different between the two measurement instances, most likely caused by remounting of a different bead sample on the stage. We attribute the difference in the estimated coma to this sample tilt, as passing a high NA emission beam through a tilted medium leads to coma that is constant over the FOV. Apparently, the aberration estimation from bead calibration data can be affected by small changes in the imaging conditions, like the sample tilt. Similar differences in imaging conditions can potentially arise between the bead calibration measurement and the actual single molecule data acquisition, implying that estimation of aberrations from the data itself not only makes the bead calibration measurement spurious, but is also inherently more reliable.

We also used the chi square goodness of fit test to assess the added value of taking the field dependency of the aberrations into account. We applied Vectorfit with all the aberrations set to zero, except for a constant amount of astigmatism (*A*_2−2_ = 87 *mλ*, equal to the average amount over the FOV estimated from the bead calibration) for estimating the axial location for the 3D data. Supplementary Figure 3 shows that the PSF model with single molecule data derived field dependent aberrations fits the data better than the model without aberrations, especially towards the edges of the FOV. A comparison of the estimated locations using the two models shows that the location difference increases with the distance from the center of the FOV (Supplementary Fig. 3d). Moreover, for the 3D data, the shift in location is different for emitters with z < 0 than for emitters with z > 0 (Supplementary Fig. 3f), which could lead to a distortion of the imaged structures when field dependent aberrations are not considered.

We compared our reconstructions to existing methods, namely Picasso for the 2D data and SMAP for the 3D data. Picasso [Picasso] uses a Gaussian PSF model where the horizontal and vertical standard deviations *σ*_*x*_ and *σ*_*y*_ are estimated for each ROI [Smith2010], whereas SMAP [SMAP] uses a spline PSF model that is uniform across the FOV, with parameters estimated from the experimental bead data. Figures 2c,d show that 2D reconstructions with Picasso are very similar to reconstructions with Vectorfit. The average localization precision, estimated by linking localizations in consecutive frames [Cnossen2020], is 10.0 nm for Vectorfit with a CRLB of 9.3 nm, better than the value of 13.5 nm found for Picasso with a CRLB of 14.2 nm. Figures 2g-j show that 3D reconstructions with SMAP are similar to reconstructions with Vectorfit, where Vectorfit shows a somewhat smaller axial spread of the localizations. The linking analysis gives here an average localization precision of 12.1 nm (*xy*) and 29.9 nm (*z*), with a CRLB of 12.1 nm (xy) and 31.1 nm (z), compared to a precision of 12.1 nm (*xy*) and 35.7 nm (*z*) for SMAP. FRC image resolution [Nieuwenhuizen2013] estimation (Supplementary Figs. 4 and 5) show a comparable performance between Vectorfit and the other used softwares.

Next, we applied our method to 3D data of NUP96 in the nuclear pore complex (NPC) in U2OS cells (FOV of about 180×180 µm) of Fu et al. [Fu2023], where the axial direction was encoded using astigmatism, and compared the outcome to that of their method FD-DeepLoc in Figure 3. The aberrations maps retrieved from the single molecule data matches well with the aberration maps obtained from the bead calibration used in FD-DeepLoc, towards the edges of the FOV, however, the bead calibration derived aberration maps show large and erratic variations, which is unlikely from the optical point of view. Figure 3b shows the experimentally found standard deviation for the estimation of the different aberrations, obtained from 30 estimations for different randomly selected subsets of 5,000 localizations, as well as their Cramer-Rao lower Bound (CRLB) values. The experimental standard deviations are on the small *mλ* scale across the whole FOV, but do not reach the CRLB values in the sub-*mλ* level. We also tested the dependence of the experimental precision on the number of localization events *M*_*s*′_ and found a scaling inversely proportional to *M*_*s*_^1/2^, in agreement with expectations (Supplementary Figs. 6 and 7). The comparison of the modelled PSF to the measured single molecule spots in 4×4 patches across the FOV, as a function of the axial position (Supplementary Video 2) shows a good match between the modelled PSF with the measured data. These validation tests indicate that our approach enables unbiased and precise aberration estimation.

**Fig. 3.**
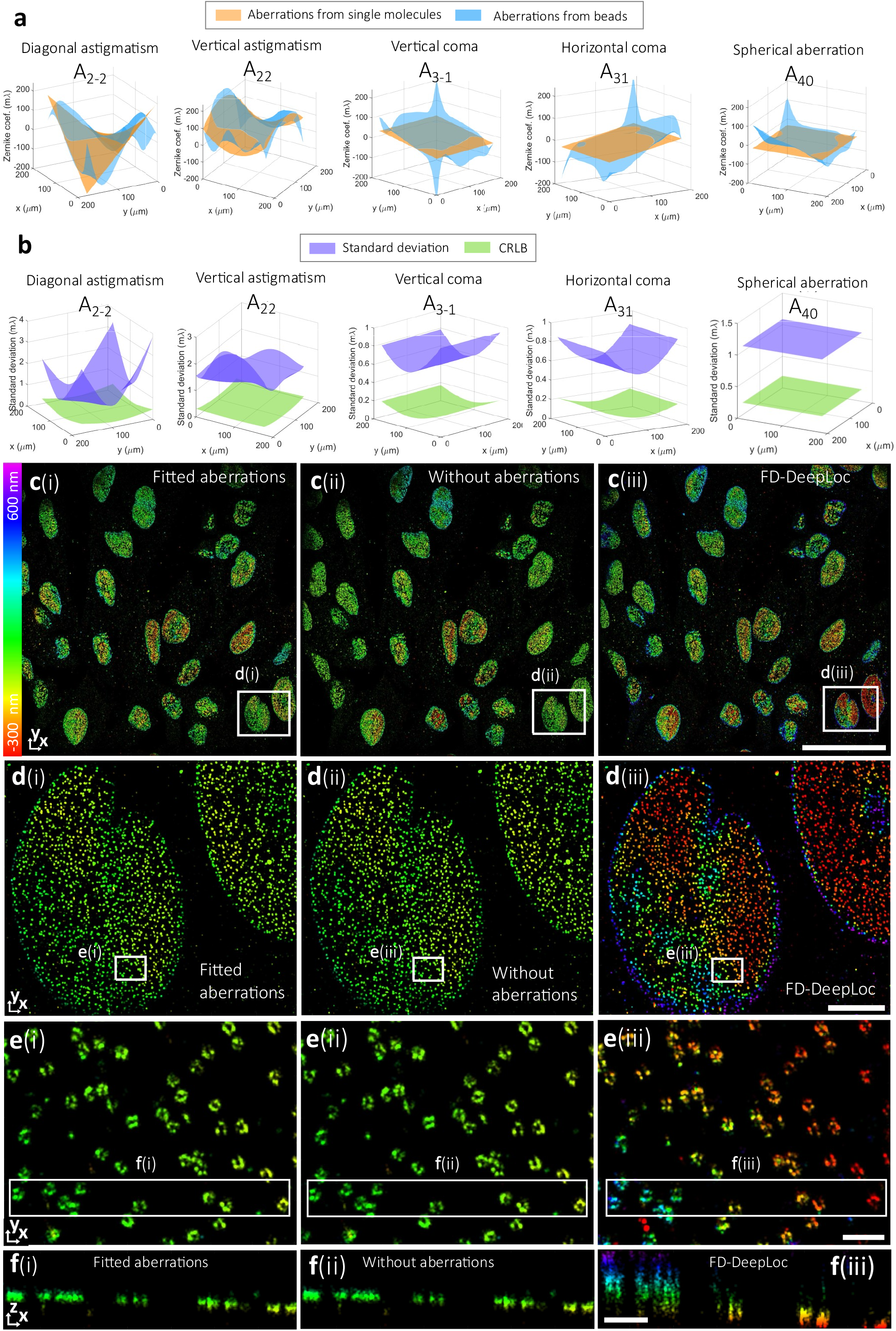
Aberration estimation and reconstructions for experimental 3D astigmatic data of NPCs. **a**, Estimated Zernike aberration surfaces from single molecules, compared to interpolated aberrations from bead z-stack calibration. **b**, Standard deviation and CRLB for the estimated Zernike coefficients. The standard deviations were calculated by repeating the estimation process 30 times with different randomly selected subsets of 5,000 localizations. **c**(i)-(iii), Full FOV images reconstructed using fitted aberrations from single molecules, without aberrations (*A*_22_ = 103 *mλ* and other Zernike modes set to zero) and FD-DeepLoc. **d**(i)-(iii), Zoomed views of the regions indicated by the boxes **d**(i)-(iii) in **c. e**(i)-(iii), Zoomed views of the regions indicated by the boxes **e**(i)-(iii) in **d. f**(i)-(iii), *x*z-cross sections of the regions indicated by the boxes **f**(i)-(iii) in **e**. Scale bars: 50 µm (**c**), 5 µm (**d**), 500 nm (**e**,**f**).

We compared the Vectorfit reconstruction with field dependent aberrations retrieved from the single molecule data to a reconstruction with Vectorfit without aberrations, were all the aberrations are set to zero, except for a constant amount of astigmatism (*A*_22_ = 103 *mλ*, in accordance with the reported rms value from the cylindrical lens) for estimating the axial location, and to the FD-DeepLoc reconstruction. In the center of the FOV, close to the optical axis, the reconstructions from the three methods are comparable (Supplementary Fig. 8). Towards the edges of the FOV, however, the aberrations become larger, leading to sizeable differences in the axial localization (see Figure 3c-f). In one corner of the FOV, the FD-DeepLoc reconstruction shows a large spread of *z*-locations within a single cell nucleus (Fig. 3d(iii)). The *xz* cross-section in Fig. 3f(iii) shows a large variation of *z*-locations in a single NPC, which seems unrealistic. In our reconstructions (Fig. 3d(i-ii) and Fig. 3f(i-ii)), the range of *z*-locations matches much better with the expected thickness of the NPC ring of around 50 nm [Wang2023, Thevasthasa2019]. At the top of the FOV we observe that Vectorfit with field dependent aberrations leads to a better reconstruction than without aberrations. (Supplementary Figure 9). A chi-square goodness of fit test (Supplementary Figure 10) shows that the Vectorfit model with field dependent aberrations outperforms Vectorfit with constant aberrations, with a chi-square ratio reaching 1.5 at the edges of the FOV. We also observe a shift in the lateral localizations up to a few tens of nanometres (Supplementary Fig. 10b) and a shift in the axial localizations of up to 150 nanometres (Supplementary Fig. 10c). The distribution of localization precision values, estimated using the method of linking localizations in consecutive frames, has a mean of 11.7 nm (*xy*) and 23.5 nm (*z*) for Vectorfit with field dependent aberrations, with a CRLB of 10.1 nm (yz) and 20.6 nm (z). For FD-Deeploc we find by the same method a localization precision of 7.9 nm (*xy*) and 23.2 nm (*z*). With Vectorfit, we achieve the CRLB and therefore attain the optimal localization precision based on the signal-to-noise ratio of the data. The axial localization precision of Vectorfit is comparable to FD-Deeploc, but the lateral localization precision is a bit worse. We speculate that the learning-based approach of FD-Deeploc could bias localizations from the same emitter, which would introduce spurious correlations in subsequent time frames, resulting in a lower measured lateral localization precision than the CRLB. Analysis of the image resolution in a grid of 4×4 patches across the FOV with FRC thus shows a slightly better image resolution for FD-DeepLoc than for VectorFit (Supplementary Figure 11).

Fully vectorial PSF fitting in a Maximum Likelihood Estimation (MLE) framework has not found its way into the practice of the SMLM field up to now. Software fitting is mostly dominated by fitting of Gaussians or spline-based models. The reason is solely that the fitting speed was too slow for the typical number of localizations (∼10^6^) in a super-resolution acquisition. CPU implementations of 2D Gaussian MLE fitting achieve currently about 10^4^ fits/s and GPU implementations even more than 10^6^ fits/s (7×7 pixel ROI). Only the phasor-based approach is equally quick on CPU [Martens2018]. Our OTF based algorithmic improvements for the fitting together with a high-end GPU (Nvidia A100) allow now 10^5^ fits/s (see Supplementary Table 1), while a good desktop GPU (Nvidia RTX A4000) has 6 · 10^4^ fits/s and a multi-core CPU still achieves 9 · 10^3^ fits/s. This brings vectorial PSF fitting times for a typical data set to less than a minute on a GPU in 2D. In 3D our algorithmic improvements are less effective (for details see Supplementary Note) which results in 6.8 · 10^3^ fits/s for a 17×17 pixel ROI on our desktop GPU which is about 30*x* slower than 3D spline-based fitting which achieves 2 · 10^5^ fits/s on a desktop GPU for a 17×17 pixel ROI [Li2024], but better than the less than 1 fit/second reported for a 17×17×41 pixel bead stack [Fu2023].

Our approach of estimating aberrations directly from single molecule data makes taking the inclusion of field dependent aberrations to the localization much more feasible because it saves doing an additional bead calibration experiment, which can also be prone to errors as shown above. The recently proposed method from Liu et al. to estimate aberrations from single molecule data can take more than half an hour on a GPU [Liu2024], while our model-based approach takes less than a minute on a CPU to estimate aberrations. In addition our work is applicable to 2D localization data while theirs is not. In 2D, however, the effect of field dependent aberrations on the localizations is less impactful than in 3D.

In future work, a user interface should be developed to make vectorial PSF fitting accessible for a broader audience. Furthermore, the NAT driven approach of estimating field dependent aberrations directly from single molecule data could be applied to other imaging modalities such as 4Pi SMLM or localization microscopy with fixed dipole emitters, and also for scanning but camera-based localization microscopy [Radmacher2024].

## Author contributions

I.D. developed the analysis model and analysed all datasets, E.S., S.H. and B.v.W. developed GPU code, S.K. and K.A.L. performed imaging of biological samples. S.S. and B.R. devised key concepts and supervised the study. S.S., B.R. and I.D. wrote the paper. All authors read and approved the manuscript.

## Acknowledgements

We thank Yutong Wang for assistance with experiments and Sheng Liu for helping with instrument modification. S.S. acknowledges funding from European Research Council (101055013) and B.R. from Nederlandse Organisatie voor Wetenschappelijk Onderzoek (17046) and from the eScience Center (NLESC.SSI.2021b.001). K.A.L and S.K. were supported by NIH grant 1R01GM140284.

## Data and code availability

Code and a software demo are available at https://gitlab.tudelft.nl/imphys/ci/vectorfit.

## Methods

### Processing pipeline

First, the set of acquired images were offset and gain corrected to convert ADUs into photon counts [Heintzmann2018, vanVliet1998a]. ROIs of size of 9 × 9 pixels for 2D data and 17 × 17 pixels for 3D data pixels are identified by a two-stage filtering process to reduce photon noise and local background followed by an intensity threshold [Huang2011, Smith2015]. In short, we apply uniform filters to the raw images with filter size 3 and 7 pixels and take the difference. We then compute the local maximum in a 7×7 pixels neighbourhood for all pixels and accept the central pixel as candidate for a single-molecule spot if its value is the local maximum and is higher than a threshold directly above the background noise level (different for each dataset). Now the 9 × 9 or 17 × 17 ROI is segmented out for all candidates and each ROI, labelled with index *s*, is extracted, leading to a total of *N*_*loc*_ candidate localization events.

A randomly selected subset of *M*_*s*_ ≪ *N*_*loc*_ of localizations is processed to determine the Legendre NAT coefficients that characterize the variation of the Zernike aberration coefficients across the FOV. To ensure an even distribution over the FOV, the localizations are selected on a 10×10 grid over the FOV where the same number of localizations is selected from each grid cell. The selected localizations are fitted using a Maximum Likelihood Estimation (MLE) algorithm with alternating local and global iteration steps. In the local iteration step the emitter position 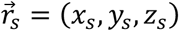, signal photon count *N*_*ph*_ and background *bg* of all *M*_*s*_ localizations are updated until they no longer improve, given the current set of NAT coefficients. For 2D data, the axial position is fitted as well, ensuring that *z*_*s*_ ≥ 0, because this leads to an improved aberration estimation compared to keeping the *z*_*s*_ value fixed. In each global update step, 20% of the *M*_*s*_ localizations are selected randomly. Then, the NAT coefficients are updated until they no longer improve, given the estimated positions, signal photon counts and background level of this subset of localization events. By using only 20% of the emitters in each step, the process is sped up significantly, while, with high probability, each of the *M*_*s*_ localization events will be used at least once in the full optimization process. The process typically converges after 6-10 steps and takes 30-45 seconds with *M*_*s*_ = 5,000 on a CPU using 12 cores (Intel Xeon w5-2455X). The NAT coefficients are randomly initialized such that the system aberration value *W*_*SYS*_ (see Supplementary Note) is equal to 40 *mλ*, because a small value different from zero appeared to result in a faster convergence. In the case of 3D data with astigmatism, an additional amount of 150 *mλ* of constant astigmatism is incorporated in the initial NAT coefficients. For 2D data, the procedure is repeated 10 times with different random initializations for the same emitters, after which the solution with the highest likelihood is selected to rule out convergence to a local maximum. This is needed to mitigate the risk of getting stuck in a local optimum during optimization. The precision of the method is estimated by repeating the process *P*_*s*_ = 30 times by taking different randomly selected subsets of *M*_*s*_ localization events. We found that several thousands of localization events are typically sufficient to reliably find the NAT coefficients (compare Supplementary Figures 6 and 7). Unless indicated otherwise, we use the NAT expansion to first order, giving 13 independent NAT coefficients that we denote by 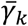 with *k* = 1,2 …, 13, taking into account the dependencies between the *γ*_*lqnm*_ as described in Supplementary Note.

For each set of NAT coefficients, there is another set of NAT coefficients that leads to the same PSF, in combination with flipping the sign of the axial coordinate *z*_*s*_. This symmetric set of coefficients is obtained by flipping the sign of the NAT coefficients that belong to Zernike coefficients *A*_*nm*_ for which *n* is even and keeping the other NAT coefficients the same. This is because flipping the sign of these even Zernike coefficients in combination with flipping the sign of *z*_*s*_ leads to the same PSF, while for the Zernike coefficients for which *n* is uneven, only flipping the sign of *z*_*s*_ leads to the same PSF. For the first-order Zernike coefficients, the even *n* modes are defocus, astigmatism and spherical aberration (*A*_20_, *A*_2−2_, *A*_22_, *A*_40_), while the odd modes are tilt and coma (*A*_11_, *A*_1−1_, *A*_3−1_, *A*_31_) where tilt is not fitted in our model because it comes down to a shift in *x*_*s*_ or *y*_*s*_. Therefore, the symmetric solution is obtained by flipping the sign of 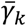 for all *k* = 1,2, …, 13, except for *k* = 10, 11,12, which are the NAT coefficients that correspond to coma. To make sure our method gives an unambiguous solution, we choose the solution for which *γ*_1_ is positive, which corresponds to the constant term of *A*_22_ (vertical astigmatism).

In a next step all *N*_*loc*_ localization events are fitted with the vectorial PSF model including the found field dependent aberrations. In these fits, as well as for the local update step in the estimation of the aberration field, we achieve major steps in efficiency by execution on GPU, as well as by several algorithmic improvements. The first algorithmic improvement is the application of phasor fitting [Maertens2018] to provide an initial estimate of the lateral position (*x*_0_, *y*_0_) of the emitter. In short, the measured photon counts *n*_*k*_ (*k* = 1,2, …, *K*^2^) of all *a* × *a* sized pixels in the ROI with pixel coordinates (*x*_*k*_, *y*_*k*_) are used to compute the four phasor components:

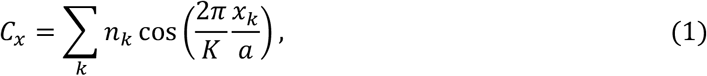

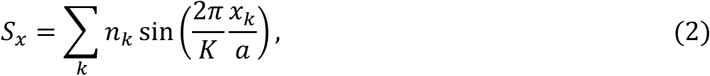

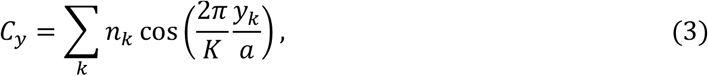

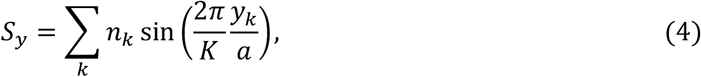

after which we find the emitter position as:

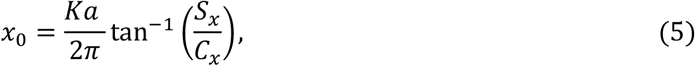

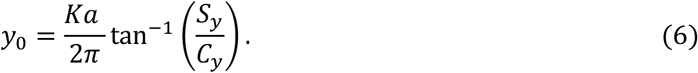

Next, a first estimate is made of the constant background photon count per pixel 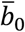 by taking the median of pixels at the rim of the ROI, and a first estimate is made for the signal photon count by 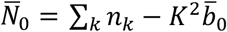. In case of 3D astigmatic fitting an initial estimate of the axial coordinate is found via computing the phasor magnitudes [Martens2018]:

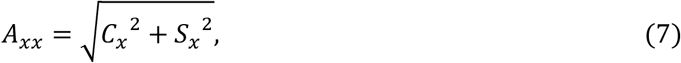

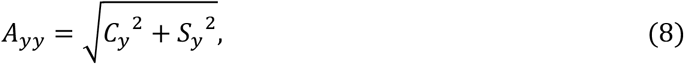

and subsequently fitting two parabolas according to [Rieger2014]:

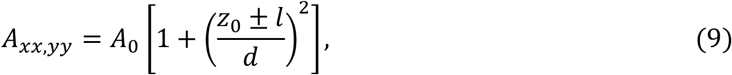

where *A*_0_ is an overall constant measuring spot width, 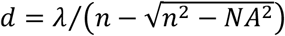 is the depth of focus and 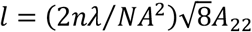 is half the distance between the two focal lines. Solving for the axial position gives:

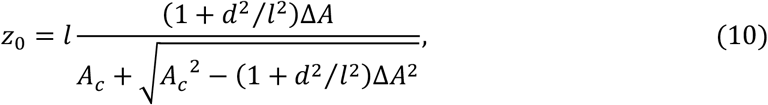

with *A*_*c*_ = *A*_*xx*_ + *A*_*yy*_ and Δ*A* = *A*_*yy*_ − *A*_*xx*_. In case the astigmatism is not vertical, we can use the more general formulas 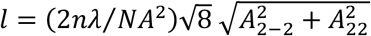 and:

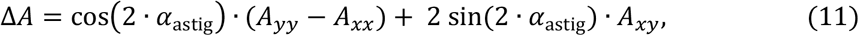

where

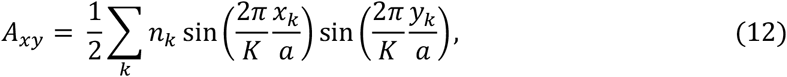

and where the astigmatism angle *α*_astig_ is determined by the amplitude of vertical and diagonal astigmatism:

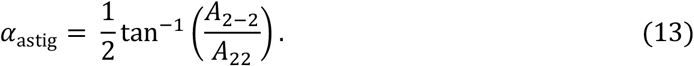

It appears that the accuracy of the initial value estimation of the axial coordinate depends on the magnitude of the astigmatism compared to size of the ROI, i.e. how well the two focal lines fit within the ROI.

The second algorithmic improvement is the use of linear regression for an improved initial estimate for signal and background photon count. To that end the vectorial PSF *H*_*k*_ = *H*(*x*_*k*_ − *x*_0_, *y*_*k*_ − *y*_0_) for 2D fitting or *H*_*k*_ = *H*(*x*_*k*_ − *x*_0_, *y*_*k*_ − *y*_0_, *z*_0_) for 3D fitting is computed (see for details below) at all pixel positions in the ROI. The improved initial values for signal photon count *N*_0_ and background photon count per pixel *b*_0_ are then found by weighted linear regression, i.e. by minimizing:

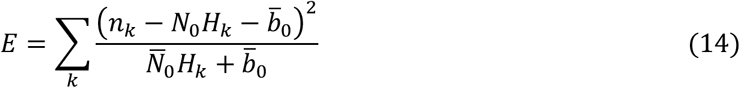

The accuracy of the initial value estimation is such that the average number of iterations needed in the final vectorial PSF localization ranges for 2D fitting from 3.7 ± 0.6 at low photon count (500 photons) down to 1.1 ± 0.3 at high photon count (10^5^ photons), and for 3D fitting from 7.1 ± 4.2 at low photon count (500 photons) down to 4.4 ± 0.8 at high photon count (10^5^ photons), as determined from a simulation study (see below for details).

The third algorithmic improvement is the use of precomputation of the Optical Transfer Function (OTF) in order to reduce the number of Fourier Transformations (FTs) per iteration. The vectorial PSF for 2D-fitting can be written as:

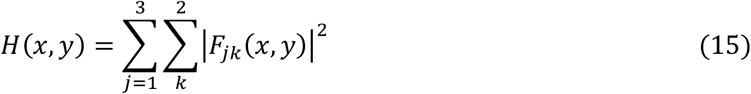

Where the matrix *F*_*jk*_(*x, y*) can be expressed as the 2D FT over the objective lens pupil:

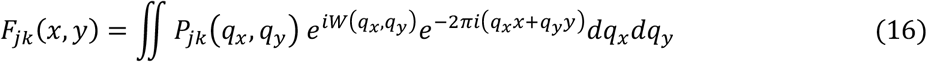

with *W*(*q*_*x*_, *q*_*y*_) the aberration function. The aberration function is computed from the NAT coefficients for the position of the center of the ROI in the FOV. Explicit expressions for the pupil matrix *P*_*jk*_(*q*_*x*_, *q*_*y*_) can be found elsewhere [Stallinga2010]. The iterative optimization in the localization algorithm requires the computation of the PSF *H*_*k*_ = *H*(*x*_*k*_ − *x*_0_, *y*_*k*_ − *y*_0_) for all pixels in the ROI, as well as the derivatives of the PSF w.r.t. the emitter coordinates. Applying the chain rule to eq. (10) implies that the derivatives of the matrix components *F*_*jk*_ (*x, y*) are then needed. These derivatives give extra factors 2*πi*(*q*_*x*_, *q*_*y*_) in the integrands for the FT. As a consequence, we need to evaluate 3 × 6 = 18 FTs in each iteration of the localization algorithm, which takes the bulk of the computation time. This can be improved by pre-computing the PSF using eq. (1), and then computing the OTF via FT:

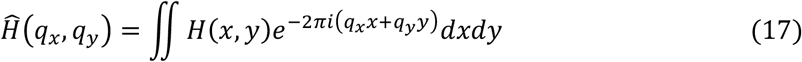

and storing the OTF in memory. For each iteration of the localization algorithm, we only need to compute the PSF and the derivatives w.r.t. *x*_0_ and *y*_0_ via the evaluation of 3 inverse FTs:

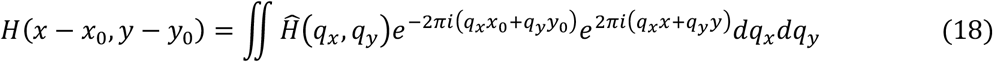

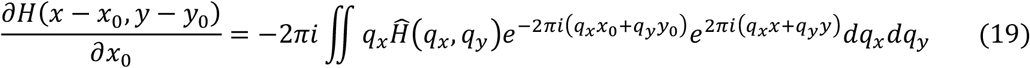

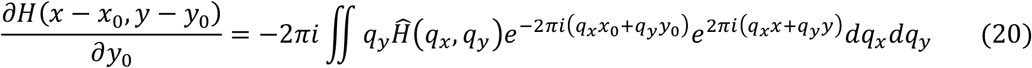

We only need to compute 3 inverse FTs now, i.e. this saves a factor of 6 in number of FTs to execute and about a 2-3*x* improvement in computation time.

For the case of 3D fitting a similar approach can be followed. Now the matrix components *F*_*jk*_ depend on *z* as well via:

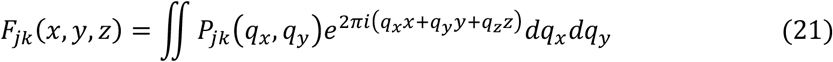

with the *z*-component of the spatial frequency vector:

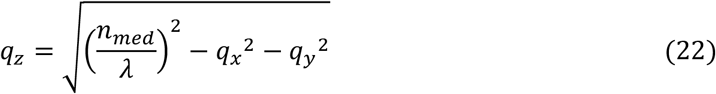

with *n*_*med*_ the refractive index of the medium and *λ* the wavelength of the fluorescence emission. Now we need to do a one-time computation of the 3D-PSF *H*(*x, y, z*), and subsequently of the 3D-OTF 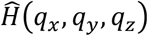 by inverse FT, and store that in memory. Note that after this inverse FT, the *z*-component of the spatial frequency vector is not restricted to be dependent on the two lateral components as in the integration over the objective pupil plane. For each iteration step in the localization algorithm, we compute the 2D-OTFs from the 3D-OTF:

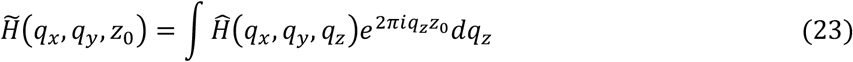

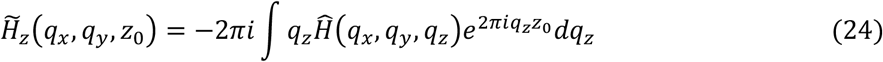

and then the PSF and the derivatives w.r.t. all the coordinates via the 4 FTs:

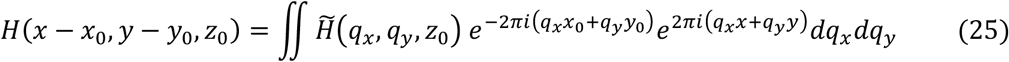

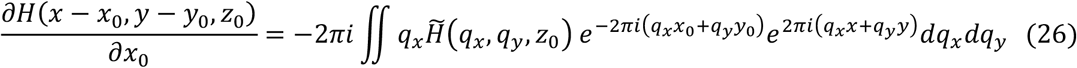

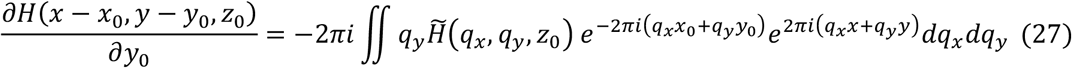

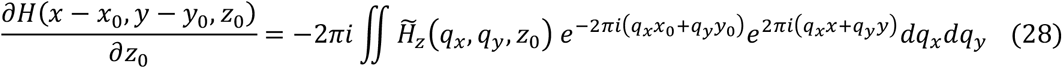

Full direct computation of the PSF and the 3 derivatives via computation of the matrix elements *F*_*jk*_(*x, y, z*) and its derivatives needs 4 × 6 = 24 FTs, so that again a factor of 6 is gained in number of FTs. The sixfold reduction of the number of FTs to execute, which make up a substantial portion of the total computational costs, leads to a 2 to 3-fold improvement in the computation time for 2D and 3D CPU and 2D GPU implementations (Supplementary Table 1). For 3D fitting on the GPU, however, no improvement is not realized. The exact cause of this remains unclear, but maybe be debit to inefficiencies in our GPU implementation of the 3D OTF computation.

In principle, the 2D or 3D OTF needs to be pre-computed for all different ROI positions in the FOV. The Zernike aberration coefficients, however, vary slowly across the FOV. In practice therefore, it is sufficient to pre-compute the OTF for 20×20 patches in the FOV. All FTs are executed with the Chirp Z-Transform (CZT) algorithm [Bakx2002], which allows for a free choice of number of samples and spacing of samples in both the spatial and the Fourier domain. For direct vectorial PSF fitting we use a pupil plane sampling with 42 × 42 for the 2D data (7 × 7) ROI and 48 × 48 for the 3D data (17 × 17) ROI. In order to perform the Fourier transforms, the pupil size *N*_*pupil*_ is chosen such that *N*_*ROI*_ + *N*_*pupil*_ − 1 is either a power of 2, or 3 times a power of 2.

For each ROI, optimization is stopped when the growth rate of the likelihood becomes smaller than a stopping tolerance *ε*_*stop*_. When a ROI has not converged after a maximum number of iterations of 10 for 2D data and 20 for 3D data, this ROI is indicated as outlier and is discarded. For experimental data we use *ε*_*stop*_ = 5 · 10^−10^, because we found that larger values of *ε*_*stop*_ lead to a worse localization precision because the optimization is stopped too early, while for smaller values, too many ROIs are incorrectly indicated as outlier. For simulation data a larger value of *ε*_*stop*_ = 10^−6^ appeared to be sufficient because with this the CRLB is already reached.

### Simulation setup

To determine the fitting speed of the vectorial PSF (Supplementary Table 1), a total of 1 million emitters were simulated using the vectorial PSF model with 2,500 photons and 10 background photons per pixel using Poisson statistics to generate noisy instances of computed single molecule spots. The wavelength was 550 nm, the numerical aperture 1.33 and the pixel size was set to *λ*/(4 · *NA*). For 2D fitting, the ROIs were fitted in batches 500,000 on a Nvidia A100 GPU, and in batches of 60,000 on other devices. The sizes for the ROIs, pupil, OTF, and the PSF for calculating the OTF were 7 × 7, 42 × 42, 42 × 42, and 129 × 129 pixels, respectively. For 3D fitting, the ROIs were fitted in batches of 20,000, and the used sizes for the ROI, pupil, OTF, and the PSF for calculating the OTF were 17 × 17, 48 × 48, 48 × 48 × 40, and 129 × 129 × 71 pixels, respectively. The simulated lateral location was randomly chosen in the central 2 × 2 pixels of the ROI. For the 3D fitting, 150 mλ of astigmatism was used in both the forward and the fitting PSF model, and the simulated axial location was randomly chosen between -500 and 500 nm. The maximum number of iterations was set to 5 for 2D fitting and 10 for 3D fitting. The computations on the Nvidia A100 GPU were performed using the DelftBlue supercomputer [DHPC2024]. To determine the number of iterations for different photon counts as described in the processing pipeline above, the average number of iterations was computed from localizing 10,000 emitters that were simulated using the vectorial PSF model with 500 and 100,000 photons. The ROI size for the 2D fitting was 9 × 9 with a pupil size of 40 × 40 and for 3D fitting the ROI size was 17 × 17 with a pupil size of 48 × 48 pixels. The other parameters were the same as described above.

### Post-processing

Sample drift was corrected on the localization data following the method of Cnossen et al. [Cnossen2021]. All images are rendered by histogram binning on a grid with super-resolution pixel size set to half the average CRLB of the localization precision, with additional Gaussian blurring with kernel size (standard deviation) equal to 2 super-resolution pixels.

For calculating the localization precision by linking of nearby localizations in subsequent frames, first the localization events with an estimated photon count between 500 and10000 and an estimated precision (CRLB) below 60 nm (*xy*) and 120 nm (*z*) were selected. Histograms of these parameters show that only a few extreme outlier events are filtered in this way. Localizations in subsequent frames closer than 60 nm (*xy*) and 120 nm (*z*), and closer than 2× the smallest estimated precision (CRLB) of the events in all 2 or 3 spatial directions were considered to originate from a single on-event. The unbiased variance over the total run length of linked localizations is considered as the measure for the experimental localization precision. This method can induce a bias for small run lengths, especially if one of the linked localizations is of low quality (e.g. the first or last event where the molecule may be on for only a part of the frame time). For that reason only longer on-events are used in the statistical evaluation (5 ≤ run length ≤ 13).

Experimental PSF data (Supplementary Video 1) was assembled by binning the localization events over 4×4 patches over the FOV and over 45 bins in the axial direction. The acquired spots for the different ROIs in each bin were 5× upsampled using Fourier space zero padding, centered at the estimated (*x*_0_, *y*_0_) position and finally summed.

### Sample preparation

To acquire the data of Figure 2, HeLa cells were plated on 25 mm coverslips in 6-well plate (Warner instruments) and allowed to adhere overnight (12-16 hours). First, cells were briefly washed with warm 1*x*PBS and then fixed using a solution containing 0.6% paraformaldehyde, 0.1% glutaraldehyde, and 0.25% Triton, all diluted in PBS for 60 seconds. This was followed by a secondary fixation using a buffer made up of 4% paraformaldehyde and 0.2% glutaraldehyde diluted in PBS for two hours. The cells were then washed twice with PBS, treated with 0.1% NaBH_4_ for 5 minutes to minimize background fluorescence due to glutaraldehyde, and further washed twice with PBS. To quench any reactive cross-linkers, the samples were incubated in 10 mM Tris for 10 minutes, followed by one PBS wash. Finally, the samples were permeabilized through a 20 minutes incubation in a solution containing 5% BSA and 0.05% Triton X-100 diluted in PBS. At the end, the samples were washed once with PBS and prepared for the labeling procedure. Cells were incubated with 15 µg/ml in volume of 350 uL of Monoclonal Anti-β-Tubulin, Mouse IgG clone AA2 (Sigma #T8328, concentration: 2mg/ml) in 2% BSA + 0.05% Triton X-100 for 5 hours. Then cells were washed once with Massive Photonics DNA-PAINT washing buffer (1×). For secondary antibody labeling, cells were incubated with 15 µg/ml in volume of 350 uL per coverslip of DBCO-DAMIG-DNA Dock-2 (Acronym of DAMIG: Donkey anti-mouse IgG) secondary antibodies (stock concentration is 500 µg/ml) in 2% BSA + 0.05% Triton X-100 for 12 hours. This is followed by 3 times wash with Massive Photonics DNA-PAINT washing buffer (1×). Before imaging, cells were washed once with Massive Photonics imaging buffer. The cell sample was placed in an Attofluor chamber then Atto655 imager-2 strands at concentration of 10 pM diluted in Massive Photonics imaging buffer was added for DNA-PAINT imaging.

### Data acquisition

To acquire the data of Figure 2, the sample was mounted on a stage of a custom-built sequential microscope [Schodt2023] controlled by custom-written software (github.com/LidkeLab/matlab-instrument-control) in MATLAB (MathWorks Inc.) [Pallikkuth2018]. A high-power 647 nm laser (2RU-VFL-P-500-647-B1R, MPB Communications) was used as an excitation laser. A 100X silicon oil immersion objective (UPLSAPO100XS, Olympus) was used to collect emitted fluorescent light. A 708/75 nm bandpass filter (FF01-708/75-25-D, Semrock) was placed in the emitted fluorescence light path. A 4f relay system consisting of 175 mm and 125 mm focal lengths lenses was set up in the emission path. To acquire the 3D data, we have set up two cylindrical lenses at the Fourier plane of focal lengths 1 m and -1 m respectively, for introducing astigmatism. The sCMOS camera (C11440-22CU, Hamamatsu) was used to detect emitted fluorescence light. At first, brightfield reference image was taken using 660 nm LED (M660L3, Thorlabs) illumination lamp and saved. During data acquisition, 100 sequences of 800 frames (a total of 80,000) were collected at 10 Hz. The saved brightfield reference image was used to realign each cell after every 4^th^ sequence of imaging [Wester2021]. The experimental settings for the 2D data are identical to those used for the 3D data, except that the pair of cylindrical lenses is removed.

---

**Supplementary Figure 1.**
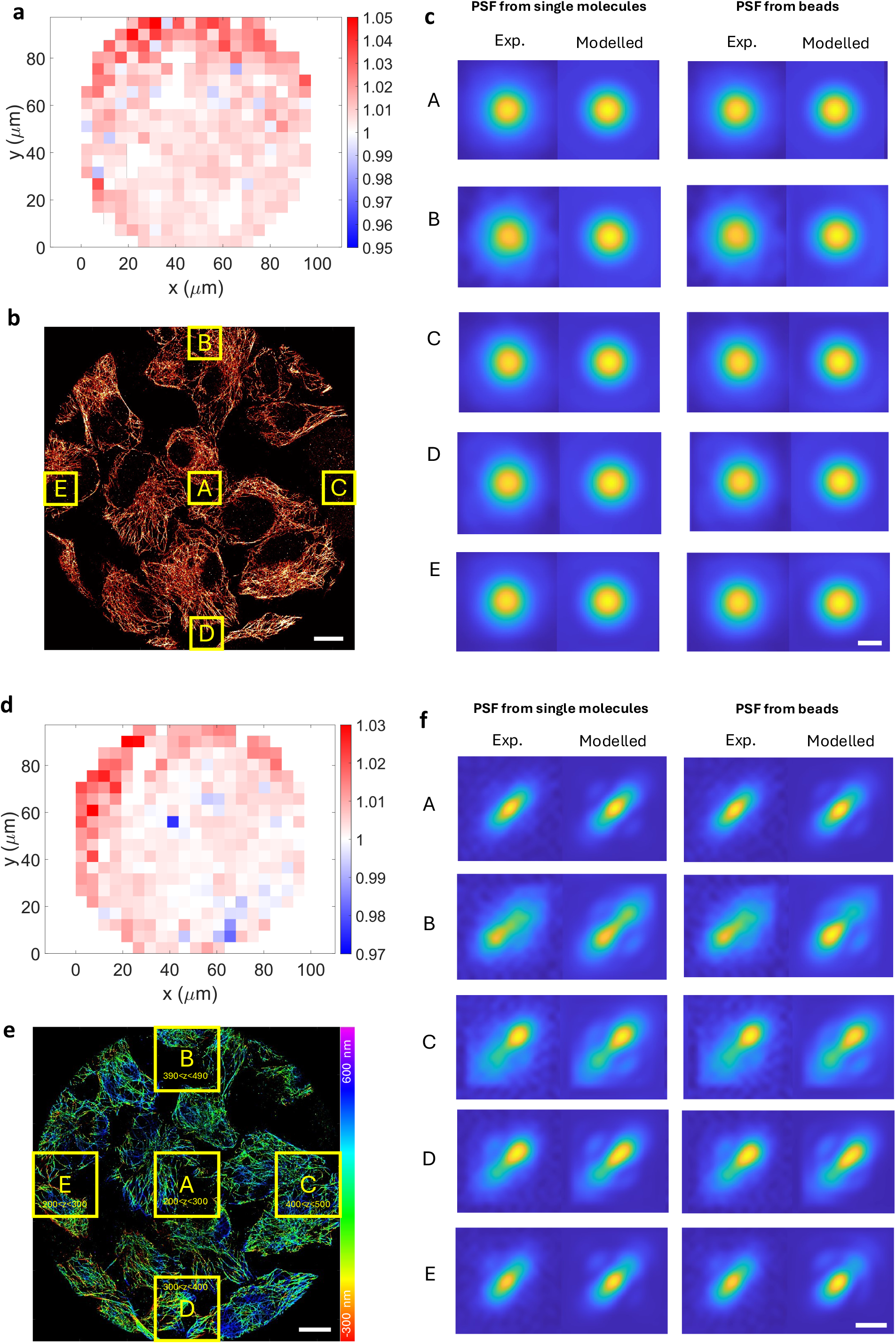
PSFs from single molecules compared to aberrations from bead calibration. **a**, The chi-squared goodness of fit (*χ*^2^ GoF) value of the PSF model with bead calibration derived aberrations to the measured ROIs divided by the *χ*^2^ GoF value of the PSF model with single molecule derived aberrations to the measured ROIs, averaged on a 20×20 tile grid over the whole FOV, for the 2D data shown in **b. b**, Full FOV 2D image indicating the 5 regions of the summed and upsampled PSFs shown in **c. c**, The average experimental and modelled PSFs within the regions as indicated in **b**, for both aberrations from single molecules and for aberrations from beads. The PSFs are obtained by centering and summing of PSFs, followed by 10-fold upsampling. Since the estimated location needed for the centering depends on the aberrations, a separate experimental PSF is shown for both the aberration choices. **d, e, f**, The same as **a, b, c**, but for the 3D data with astigmatism. Scale bars, 10 μm (**b**,**e**) 200 nm (**c**), 500 nm (**f**).

**Supplementary Figure 2.**
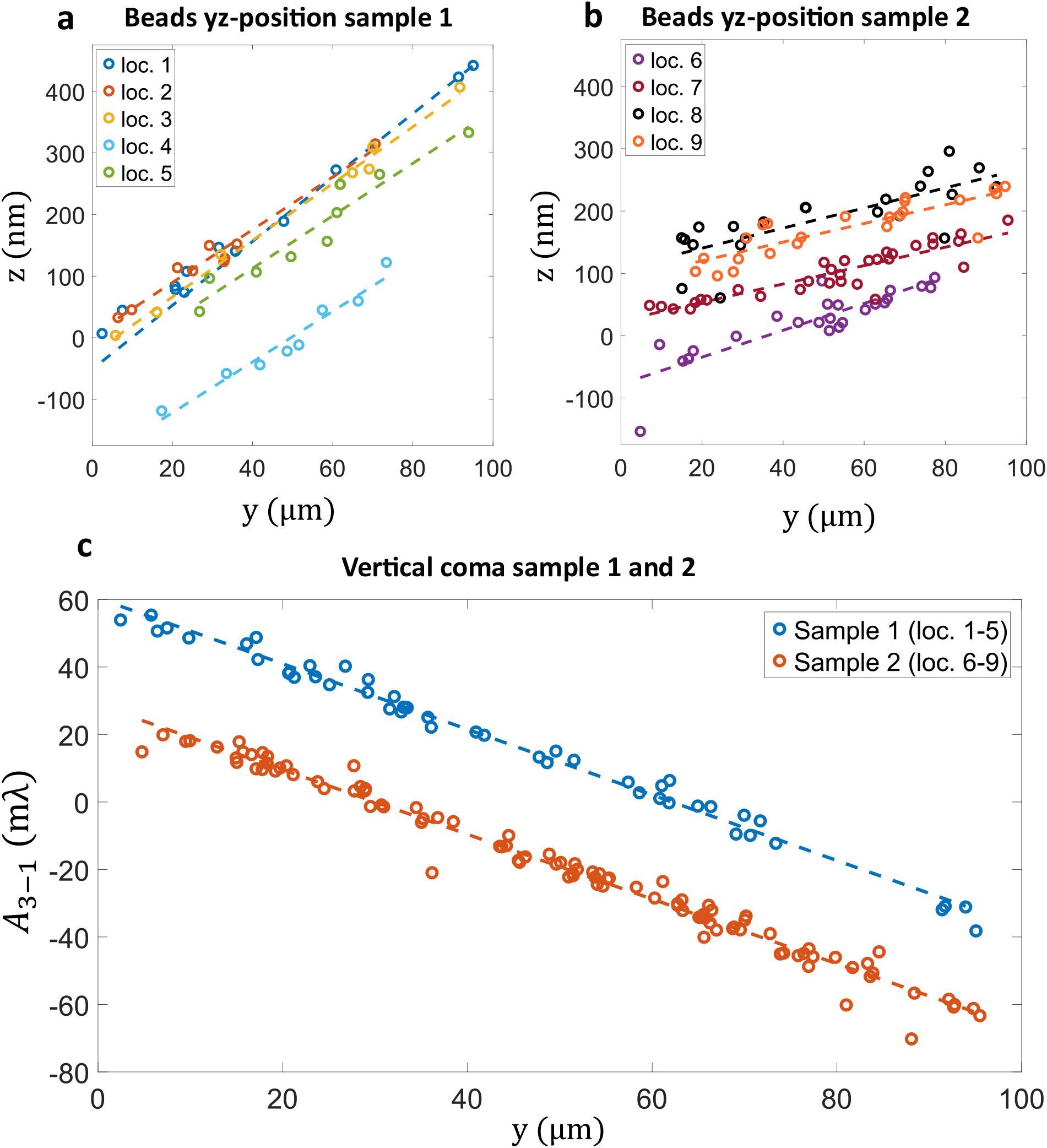
Aberration calibration with beads on two different samples. Results of two bead calibration with two different bead samples, performed on two consecutive days. From sample 1, a z-stack was recorded at 5 different locations (loc. 1-5) and from sample 2, a z-stack was recorded at 4 different locations (loc. 6-9). **a**, Estimated yz-positions of sample 1 for each location. **b**, Estimated yz-positions of sample 2 for each location. **c**, Estimated Zernike coefficient *A*_3−1_ (vertical coma) for sample 1 and sample 2.

**Supplementary Figure 3.**
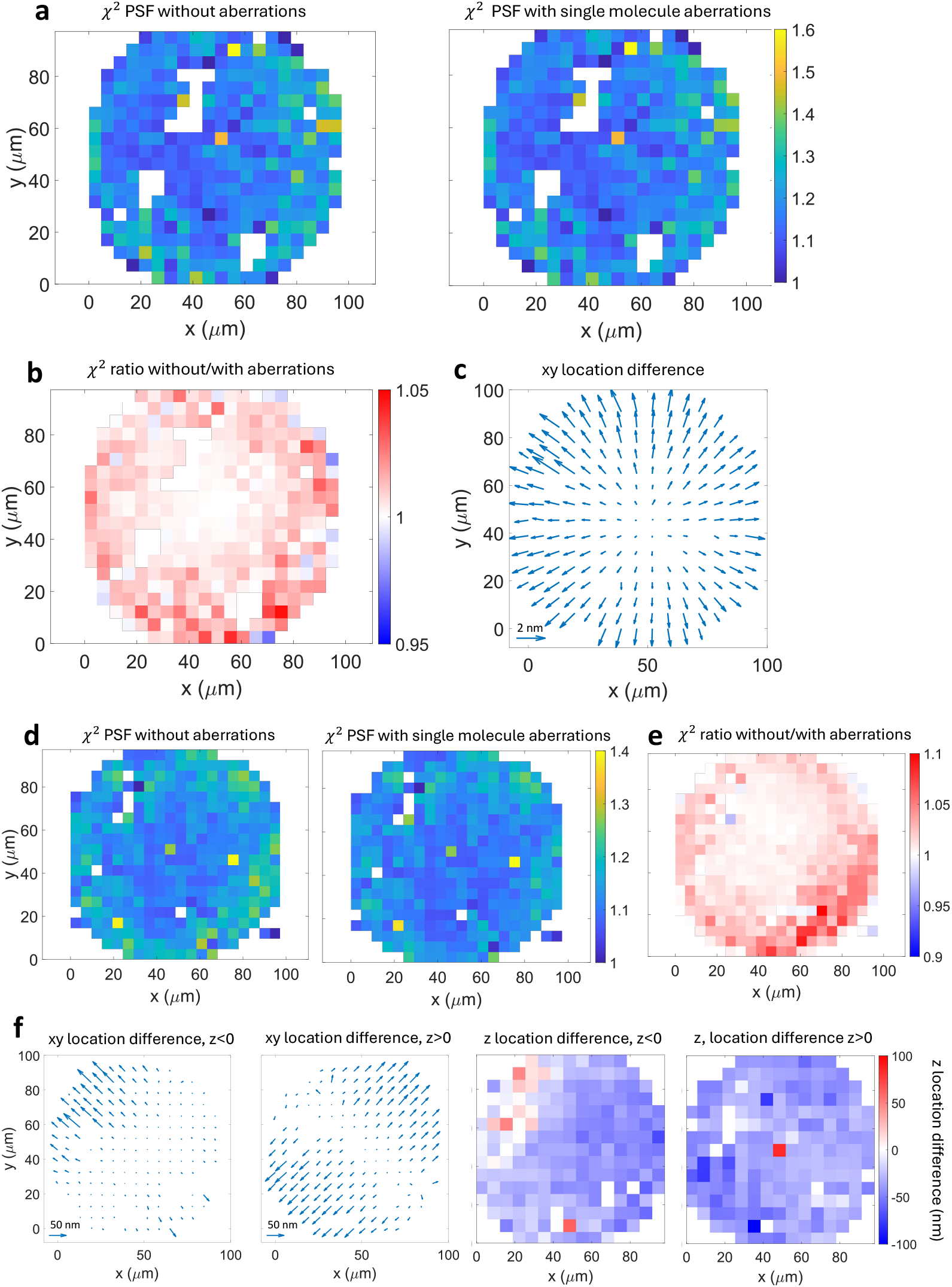
Comparison of goodness-of-fit and locations with and without aberrations. **a**, The chi-square goodness of fit value of the PSF model to the measured ROIs, averaged in 20 × 20 patched over the FOV, for the PSF model without aberrations, and the PSF model with field dependent aberrations from single molecules, for the 2D data. The values are normalized by the theoretically expected value for shot noise statistics. **b**, The ratio of the chi-square value without aberrations to the chi-square value with single molecules aberrations from **a. c**, The difference in the estimated xy locations that result from the PSF model without aberrations and the PSF model with single molecules aberrations, averaged over 15 × 15 patches over the FOV. **d-f**, show the same as **a-c**, but for the 3D data, where the xy and z location differences in **f** are broken down into emitters below (*z* ≤ 0) and above the focal plane (*z* ≤ 0).

**Supplementary Figure 4.**
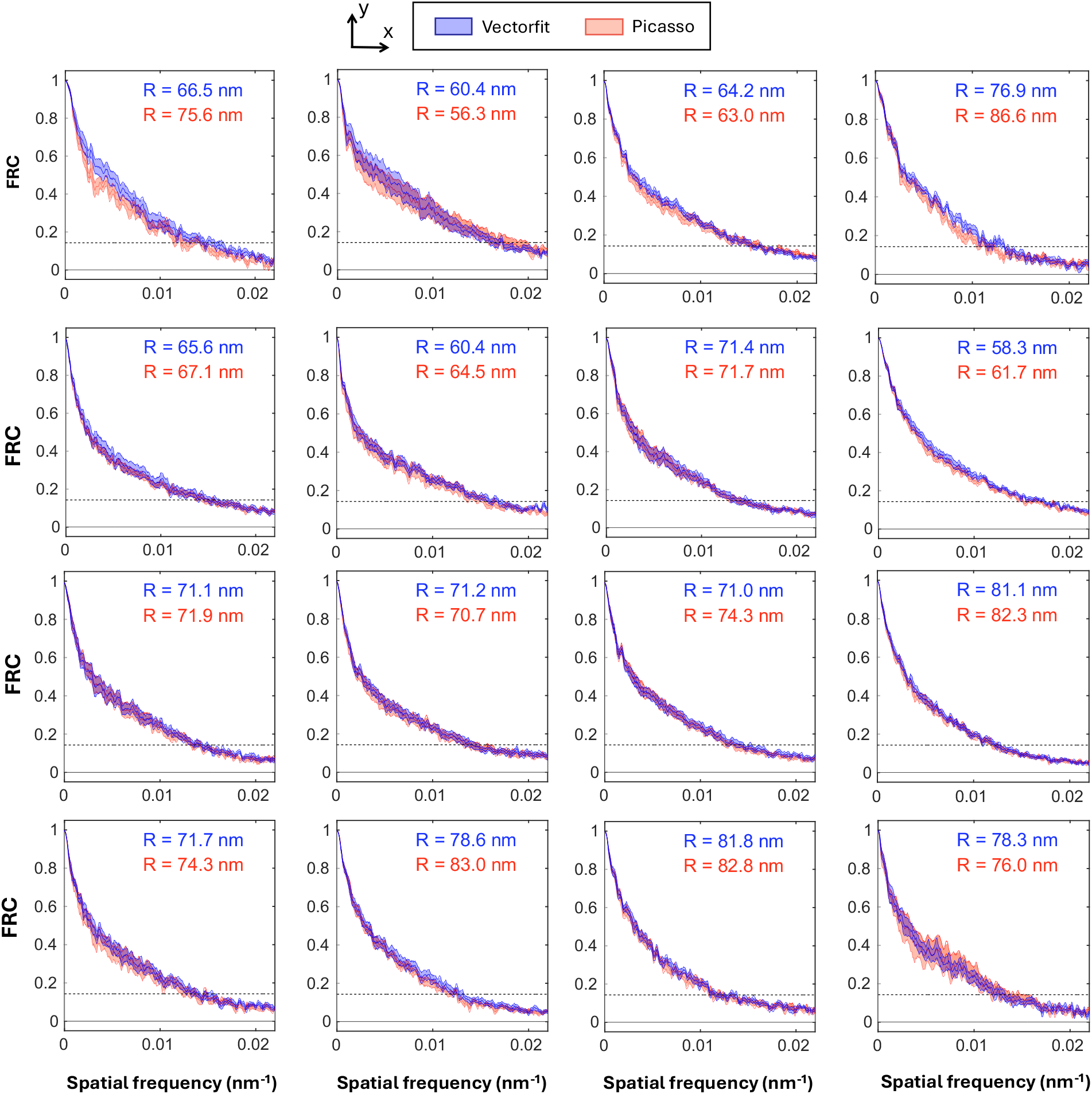
FRC analysis of 2D data of microtubili. FRC curves on a 4×4 grid of the FOV for localization data from Vectorfit and Picasso (Gaussian). The FRC analysis was done after linking localizations in consecutive frames and filtering out the brightest 10^−6^ fraction of super-resolution pixels, to eliminate hot pixels or persistently active emitters that could bias the computed resolution. Both the mean FRC curves and the spread quantified by the standard deviation over 10 repetitions are plotted. The FRC resolutions are shown in each subplot. The standard deviations are in the order of a few nm for all resolution values.

**Supplementary Figure 5.**
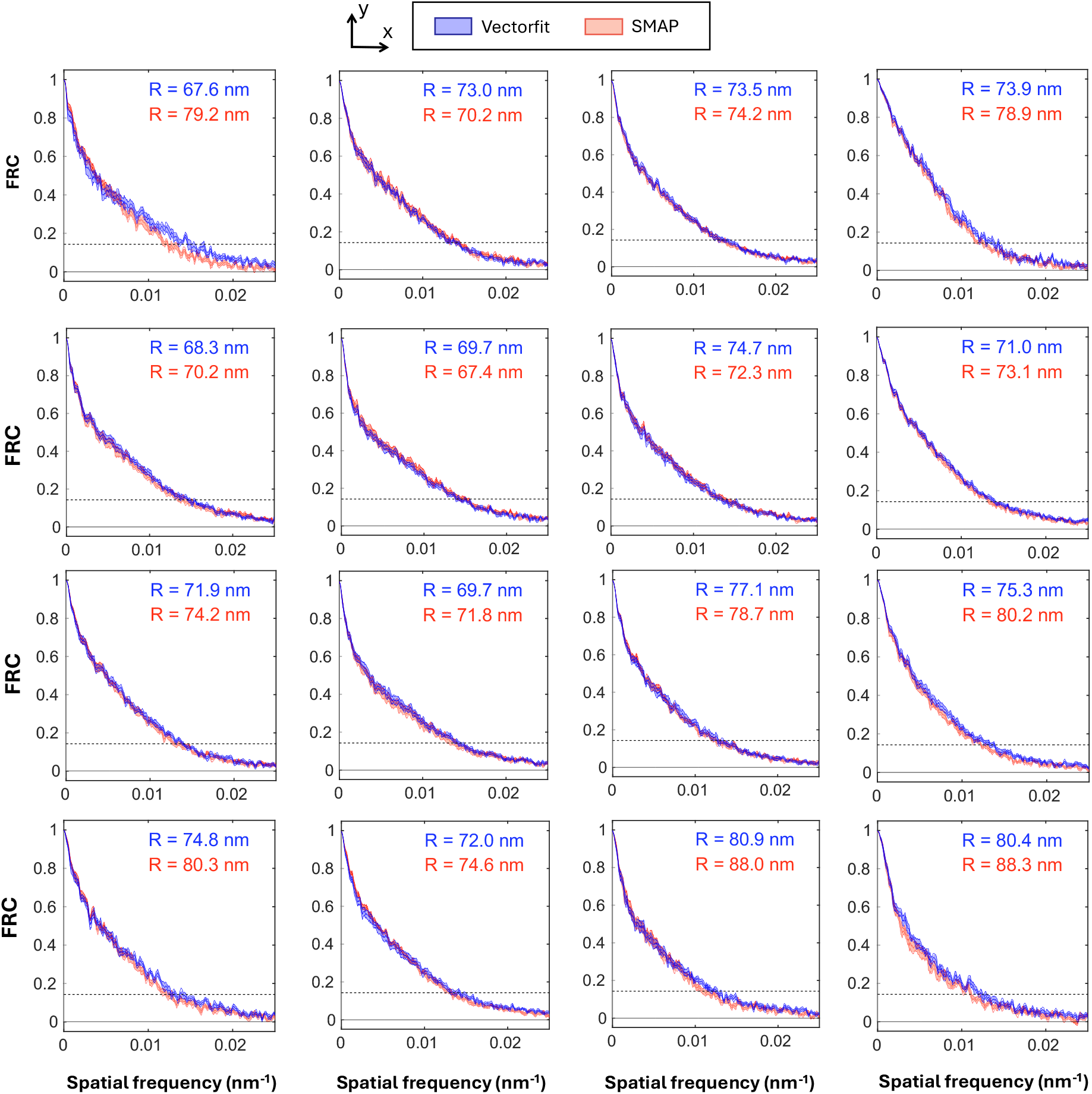
FRC analysis of 3D data of microtubili. FRC curves on a 4×4 grid of the FOV for localization data from Vectorfit and SMAP (spline). The FRC analysis was done after linking localizations in consecutive frames and filtering out the brightest 2 · 10^−5^ fraction of super-resolution pixels, to eliminate hot pixels or persistently active emitters that could bias the computed resolution. Both the mean FRC curves and the spread quantified by the standard deviation over 10 repetitions are plotted. The FRC resolutions are shown in each subplot. The standard deviations are in the order of a few nm for all resolution values.

**Supplementary Figure 6.**
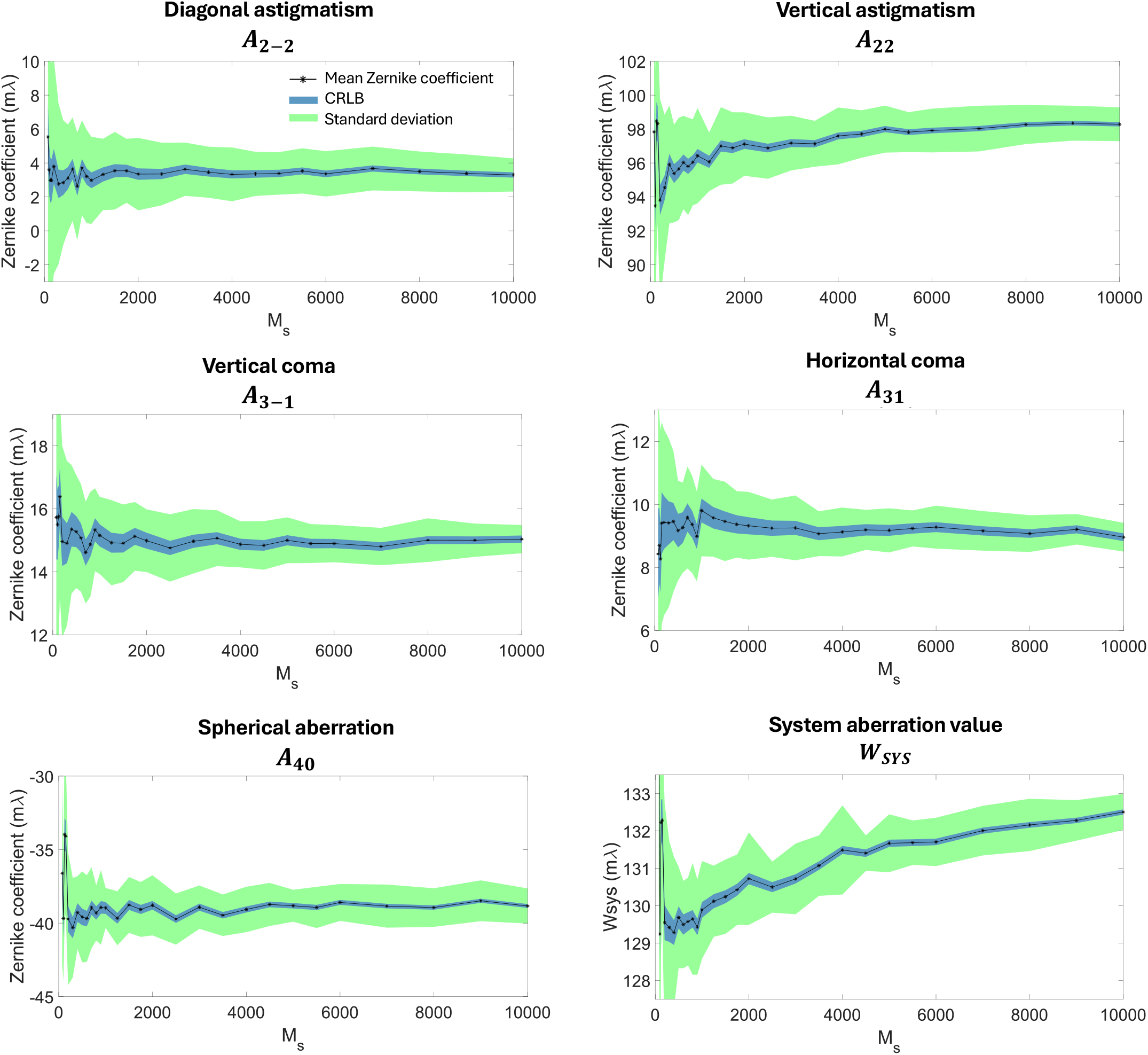
Estimated aberrations as a function of the number of spots *M*_*s*_. The standard deviation of the estimated Zernike coefficients was calculated by repeating the estimation 30 times with different randomly selected subsets of *M*_*s*_ localization events. For each value of *M*_*s*_, the Zernike coefficient CRLB was calculated. The Zernike coefficients, standard deviations and the Zernike coefficient CRLBs are averaged over the FOV. The bottom right panel shows the system aberration level (see Supplementary Note).. The 3D data of NPCs from Fig. 3 was used as input here.

**Supplementary Figure 7.**
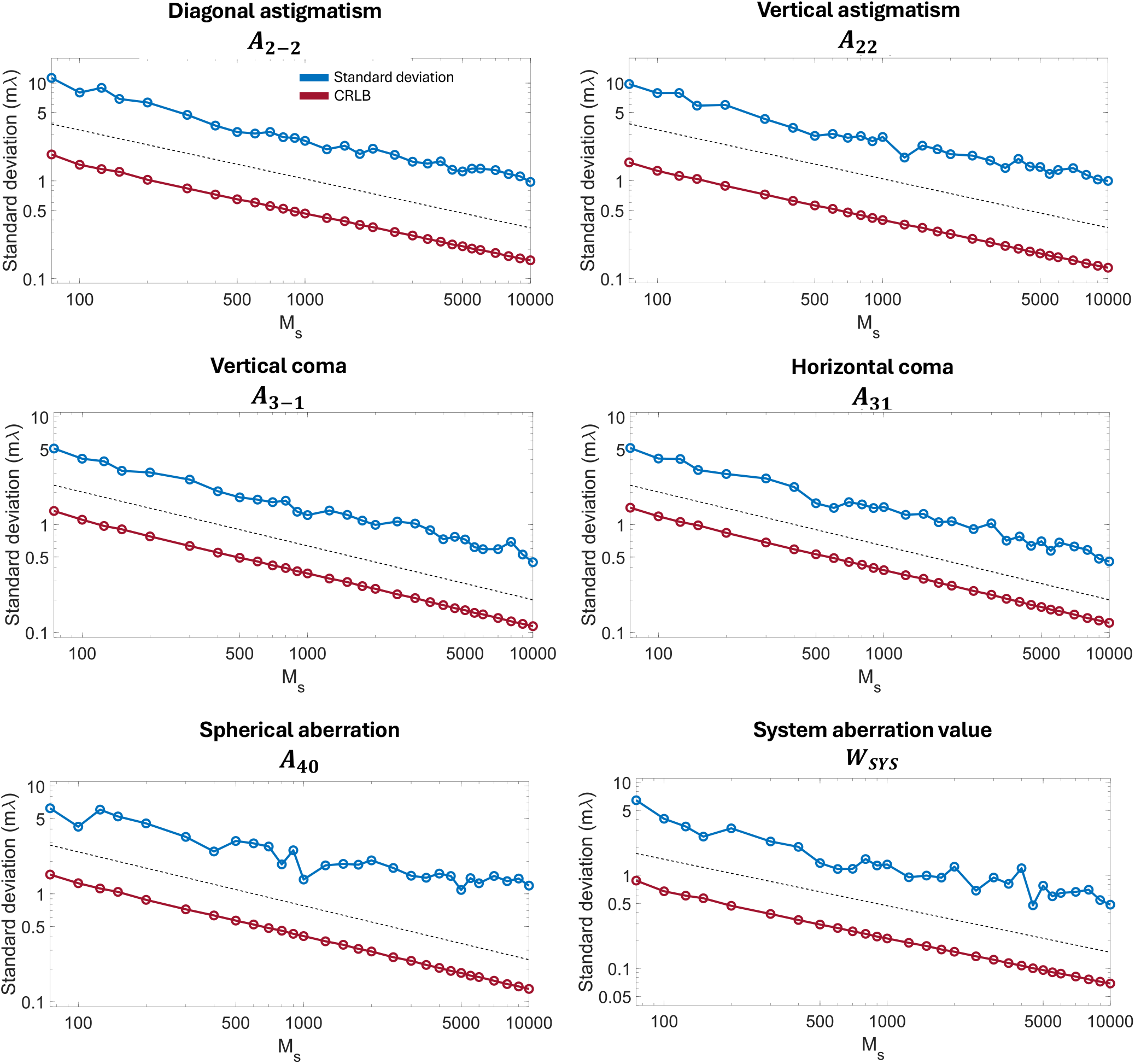
Log-log plot of the precision of the Zernike coefficient estimation as a function of *M*_*s*_ compared to the CRLB. The standard deviation of the estimated Zernike coefficients was calculated by repeating the method 30 times with different randomly selected subsets of *M*_*s*_ localization events. For each value of *M*_*s*_, the Zernike coefficient CRLB was calculated. The standard deviations and the Zernike coefficient CRLBs are averaged over the FOV. The dotted line has a slope of -0.5. The bottom right panel shows the standard deviation and CRLB of the system aberration level (see Supplementary Note). The 3D data of NPCs from Fig. 3 was used as input here.

**Supplementary Figure 8.**
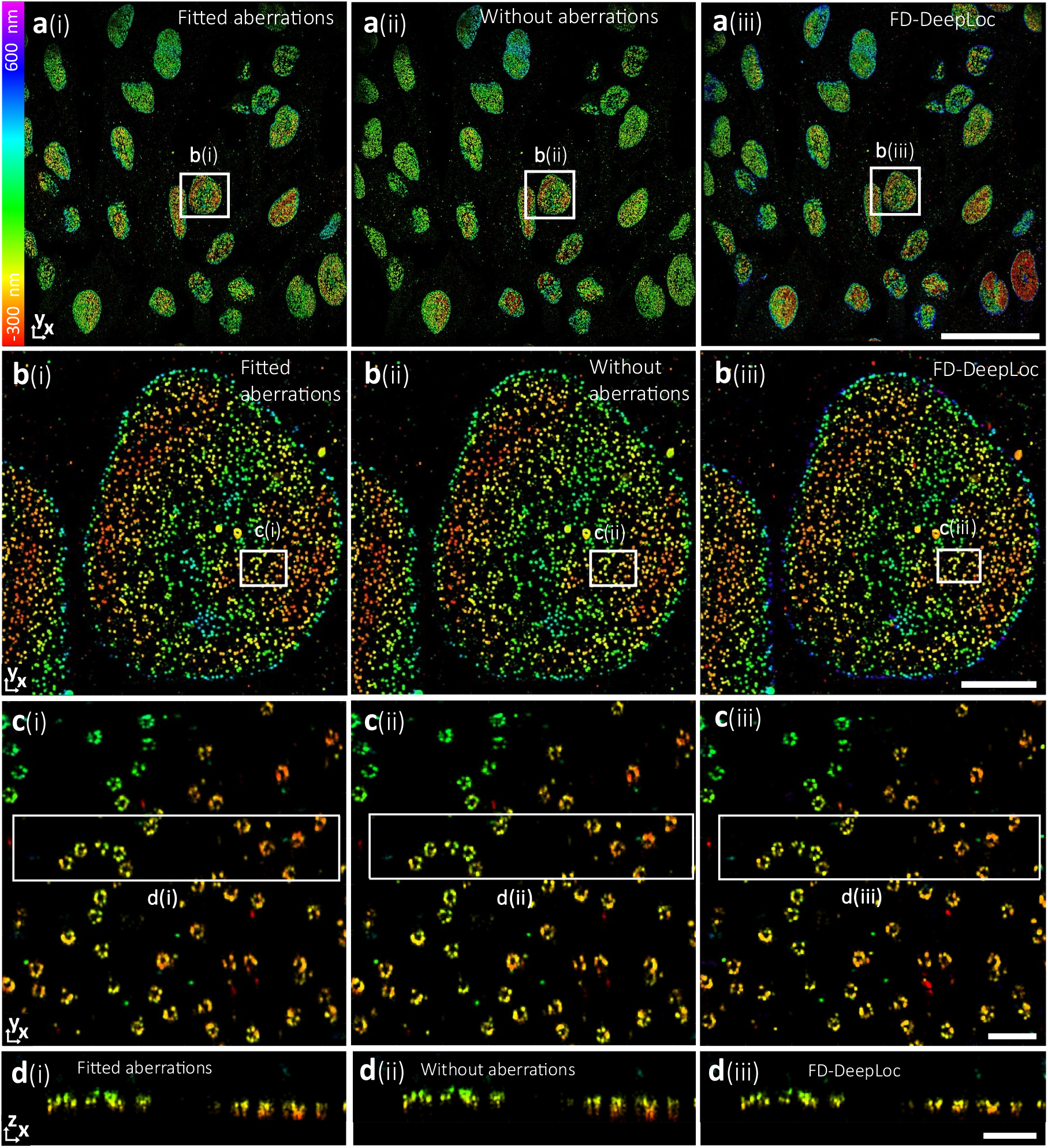
Reconstructions for experimental 3D astigmatic data of NPCs in the center of the FOV. **a**(i)-(iii), Full FOV images reconstructed with estimated aberrations from single molecules, without aberrations (*A*_22_ = 103 *mλ* and other Zernike modes set to zero), and FD-DeepLoc, **b**(i)-(iii), Zoomed views of the regions indicated by the boxes b(i)-(iii) in **a. c**(i)-(iii), Zoomed views of the regions indicated by the boxes c(i)-(iii) in **b. d**(i)-(iii), *x*z-cross sections of the regions indicated by the boxes d(i)-(iii) in **c**. Scale bars, 50 µm (**a**), 5 µm (**b**), 500 nm (**c**,**d**).

**Supplementary Figure 9.**
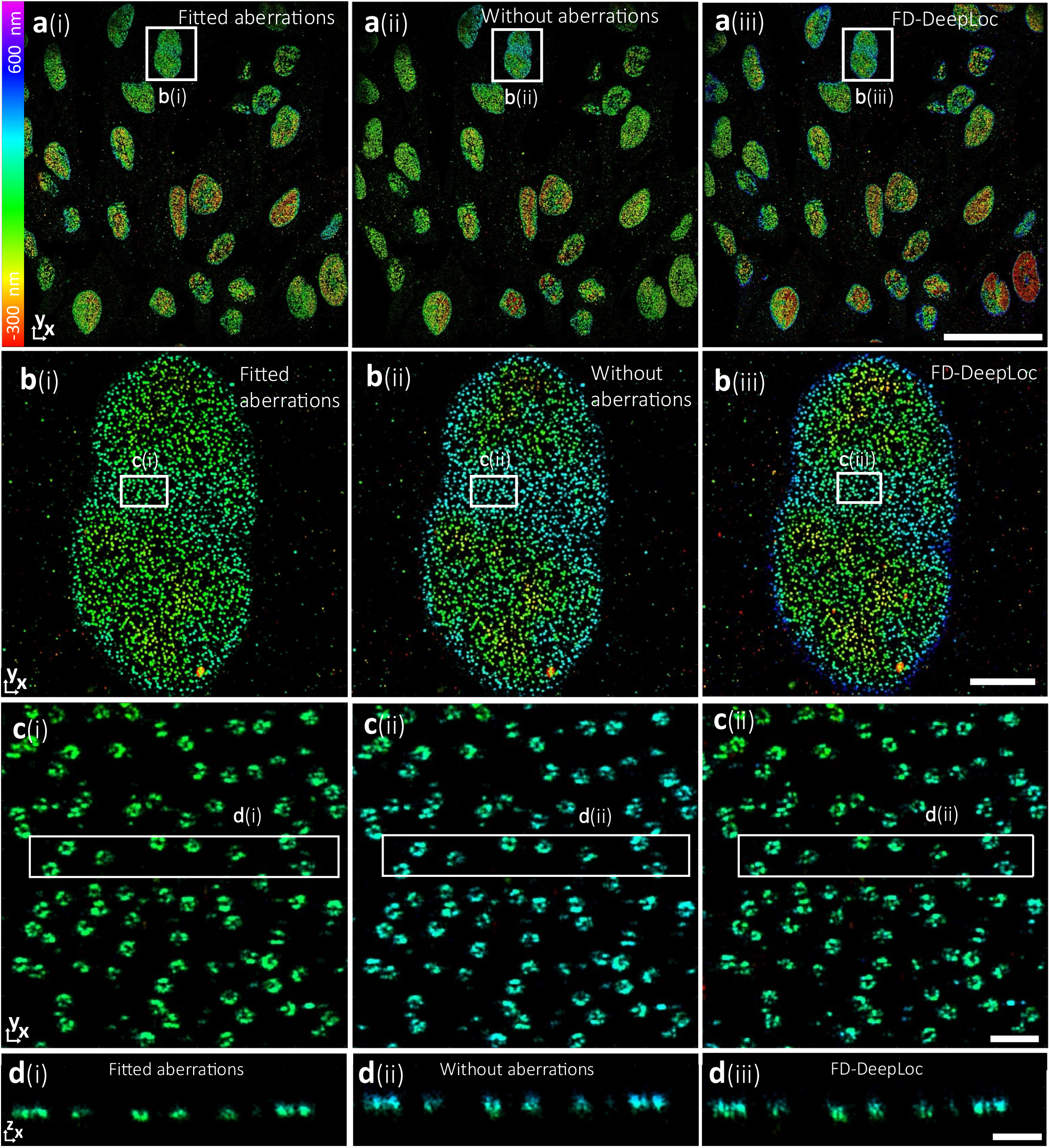
Reconstructions for experimental 3D astigmatic data of NPCs at the top of the FOV. **a**(i)-(iii), Full FOV images reconstructed with fitted aberrations from single molecules, without aberrations (*A*_22_ = 103 *mλ* and other Zernike modes set to zero), and FD-DeepLoc, **b**(i)-(iii), Zoomed views of the regions indicated by the boxes B(i)-(iii) in **a. c**(i)-(iii), Zoomed views of the regions indicated by the boxes **c**(i)-(iii) in **b. d**(i)-(iii), *x*z-cross sections of the regions indicated by the boxes **d**(i)-(iii) in **c**. Scale bars: 50 µm (**a**), 5 µm (**b**), 500 nm (**c**,**d**).

**Supplementary Figure 10.**
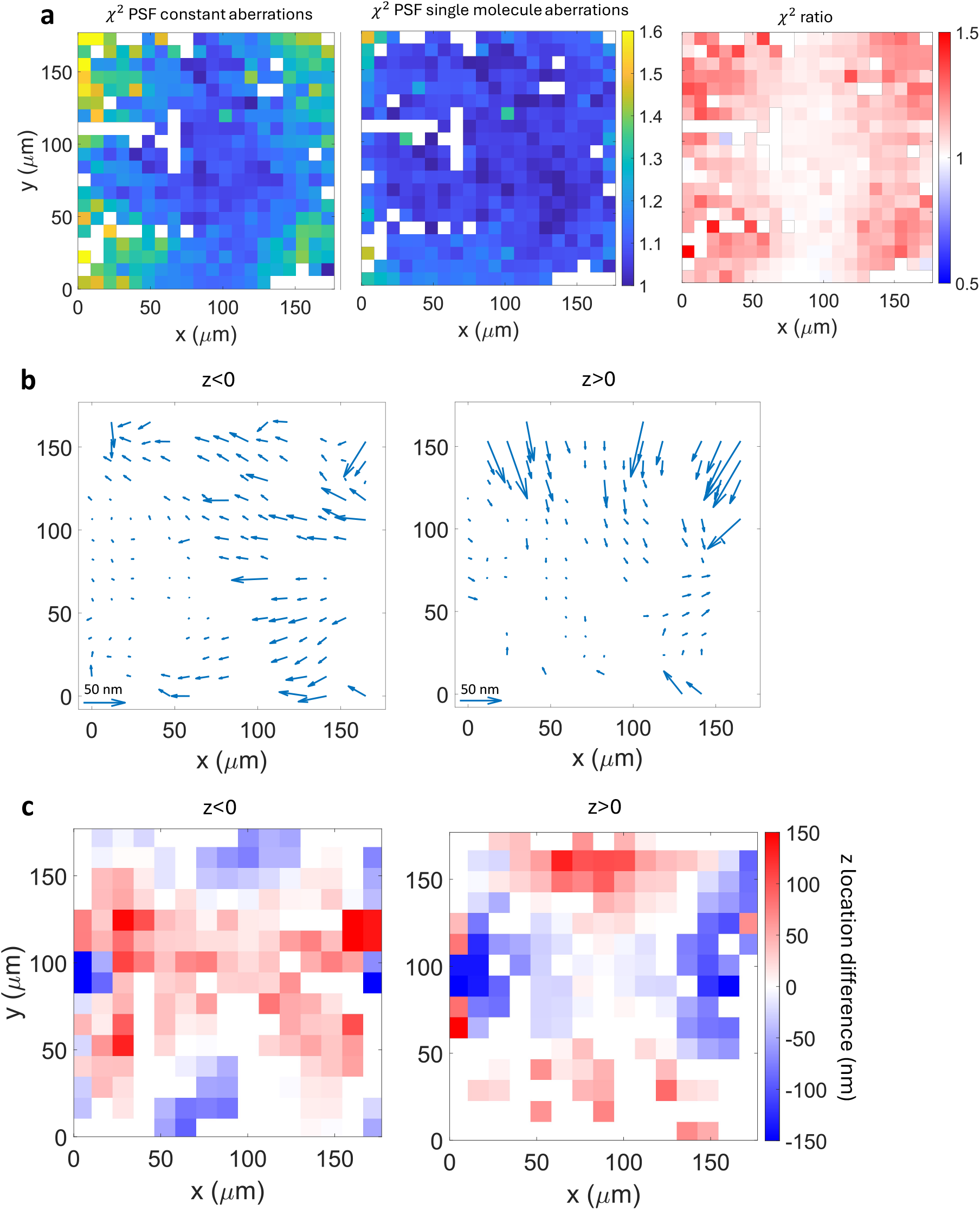
Comparison of goodness-of-fit and locations with and without aberrations. **a**, The chi-square goodness of fit value of the PSF model to the measured ROIs, averaged in 20 × 20 patched over the FOV, for the PSF model without aberrations (*A*_22_ = 103 *mλ* and other Zernike modes set to zero), and the PSF model with field dependent aberrations from single molecule data, and the ratio of the chi-square values without and with aberrations. The chi-square values are normalized by the theoretically expected value for shot noise statistics. **b**,**c**, The difference in the estimated xy-locations (**b**) and the z-locations (**c**) that result from the PSF model without aberrations and the PSF model with single molecules aberrations, averaged over 15 × 15 patches over the FOV, broken down

**Supplementary Figure 11.**
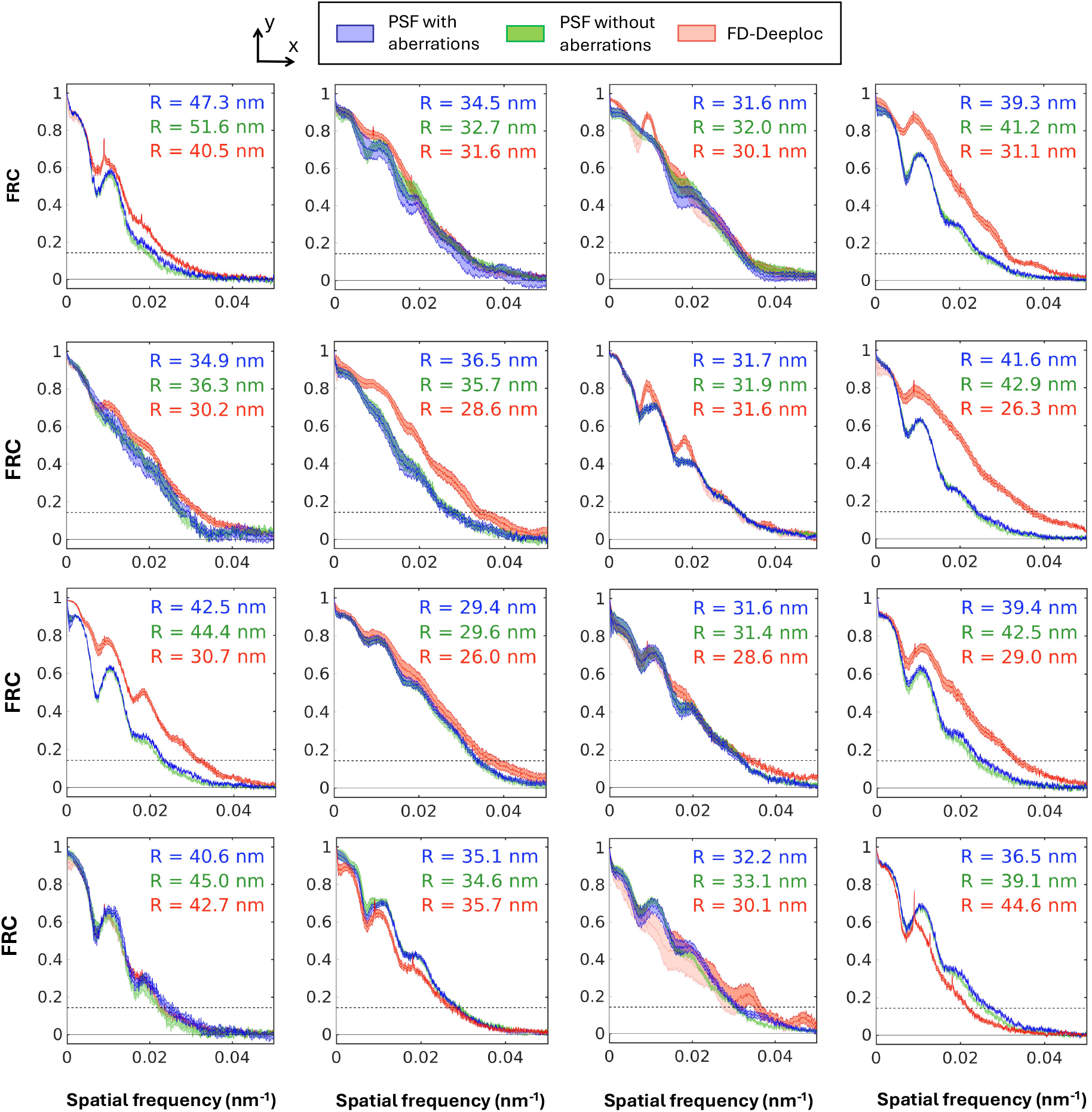
FRC analysis of 3D data of NUP96 in the NPC. FRC curves on a 4×4 grid of the FOV for localization data from Vectorfit with aberrations from single molecules, from Vectorfit without aberrations (with only 103 mλ constant astigmatism) and from FD-Deeploc. The FRC analysis was done after linking localizations in consecutive frames and filtering out the brightest 10^−6^ fraction of super-resolution pixels, to eliminate hot pixels or persistently active emitters that could bias the computed resolution. Both the mean FRC curves and the spread quantified by the standard deviation over 10 repetitions are plotted. The curves are not smooth, but have peaks that are most likely due to the dominant spatial frequency of the NPC structure (diameter around 100 nm), with first and second harmonic giving visible peaks. The FRC resolutions are shown in each subplot. The standard deviations are in the order of a few nm for all resolution values.

## Supplementary Note

### Implementation Nodal Aberration Theory (NAT) with Legendre polynomials

The key idea of NAT is to develop the Zernike aberration coefficients in a Taylor series of low order polynomials of the coordinates in the Field Of View (FOV), with the benefit of symmetry relations that reduce the number of independent coefficients. This applies to optical imaging systems with small field angles, such as microscopes or telescopes. In the following we will derive such Taylor series for 1st and 2nd order aberration theory, following the pioneering work of Shack and Thompson [Shack1980, Thompson2005] and Tessieres [Tessieres2003]. We will add a new principle to NAT, namely the use of expansions in Legendre polynomials to replace the plain Taylor series. Such an orthogonal basis function set is better adapted to the square shape of the FOV in camera based imaging, and enables a straightforward measure for the average aberration performance across the FOV, the so-called system aberration level.

#### Legendre polynomial expansion

In order to keep track of the typical or average performance across the FOV it is useful to know the system aberration value from:

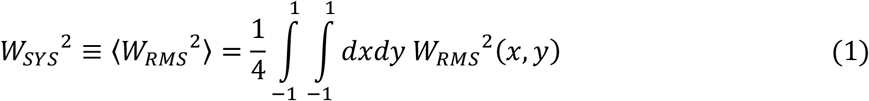

where *x* and *y* are scaled coordinates in the FOV of size *L*_*x*_ × *L*_*y*_:

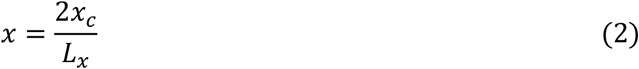

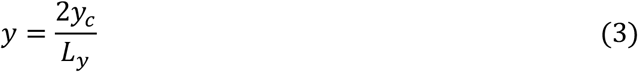

with *x*_*c*_ and *y*_*c*_ the physical FOV coordinates. The RMS aberration value at point (*x, y*) in the FOV is computed from the sum over the squares of the Zernike coefficients:

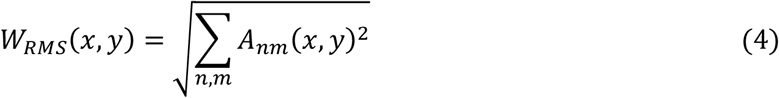

The definition of the system RMS aberration value suggests that an expansion of the Zernike aberration coefficients *A*_*nm*_(*x, y*) in orthogonal functions may be useful. This has been done in terms of Zernike modes for circularly shaped FOVs [Kwee1993]. In typical imaging systems, however, we have a rectangular, often square, FOV, defined by the camera. This implies that expanding in a complete set of functions that are orthogonal on the unit square is more practical.

Such an expansion is possible using Legendre polynomials *P*_*n*_(*x*), which are a complete set of orthogonal functions on the interval [−1,1]. Starting from the lowest orders *P*_0_(*x*) = 1 and *P*_1_(*x*) = *x*, they can be easily computed using the recurrence relation:

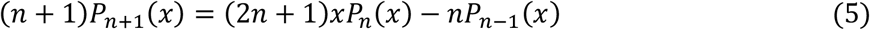

This results in *P*_2_(*x*) = (3*x*^2^ − 1)/2, *P*_3_(*x*) = (5*x*^3^ − 3*x*)/2, … They satisfy the orthogonality relation:

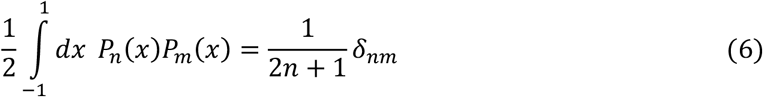

with *δ*_*nm*_ the Kronecker delta. Legendre polynomial products can be used as an orthogonal and complete set of functions on the unit square:

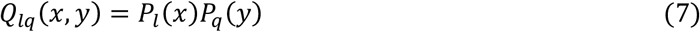

The expansion of the Zernike coefficients across the FOV can now be written as:

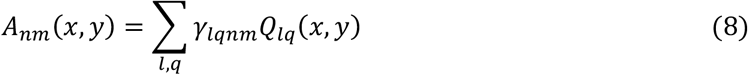

where the expansion coefficients *γ*_*lqnm*_ are henceforth called the Legendre NAT coefficients. The system RMS aberration follows from the orthogonality of the Legendre polynomials as:

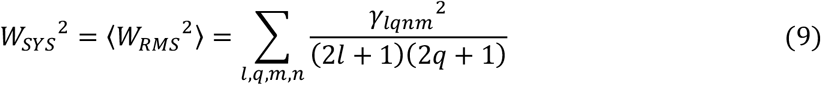

This enables a quantitative comparison of the importance of the Legendre NAT coefficients, both in absolute terms (compared to the Maréchal’s diffraction limit) and in relative terms (compared to the whole set of Legendre NAT coefficients).

#### NAT for 1st order aberration theory

Aberration theory including the impact of misalignment is developed using the pupil coordinates 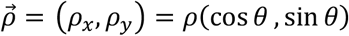, the field coordinates 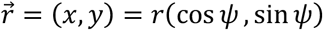, and misalignment vectors of the form 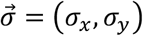 that describe lateral displacements from the aligned position. These can form 5 different rotational invariants that depend on 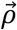 and/or 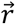, namely the inner products 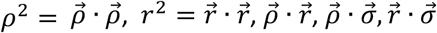. The aberration function is expanded in powers of these 5 invariants with coefficients *α*_*pqsuv*_:

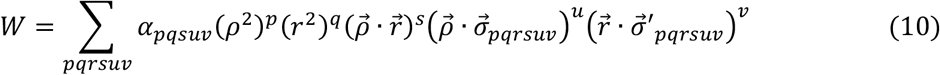

where the misalignment vectors 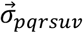 and 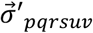 are in principle different for each term in the series expansion. The expansion order is defined using the powers of 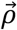 and 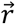 and 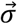 combined. For order *j* we keep the terms that satisfy:

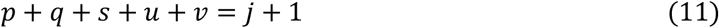

This implies that the radial order *n* = 2*p* + *s* + *u* and the azimuthal order *m* = *s* + *u* satisfy:

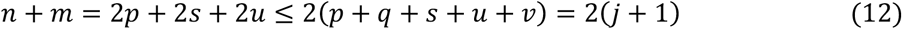

The combinatorial problem of finding integer values {*p, q, s, u, v*} that satisfy Eq. (11) is the same as dividing *j* + 1 identical balls over *k* = 5 ordered bins. There are:

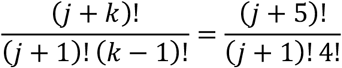

different terms (5, 15, 35, 140, … terms for *j* = 0, 1, 2, 3, …). Of these terms we can ignore the piston terms with radial order 2*p* + *s* + *u* = 0. Then *p* = *s* = *u* = 0 and the combinatorial problem boils down to distributing *j* + 1 identical balls over *k* = 2 unordered bins, which has *j* + 2 different possibilities. We can also ignore the terms linear in 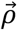, corresponding to tip/tilt that possibly depends on field coordinates, i.e. distortion. As distortion cannot be assessed without a reference object, which is usually absent, we will also ignore these terms. Now 2*p* + *s* + *u* = 1 (terms linear in *ρ*), which has as solution *p* = 0 and either *s* = 0 and *u* = 1 or *s* = 1 and *u* = 0. There are 2(*j* + 1) of these terms. These restrictions reduce the number of independent terms to 1, 8, 25, 127, … for *j* = 0, 1, 2, 3, … The number of independent parameters associated with these invariants is 1 if *u* = *v* = 0 (the pre-factor *α*_*pqsuv*_), 2 if *u* > 0 and *v* = 0 (the misalignment vector components *σ*_*x*_ and *σ*_*y*_) or *u* = 0 and *v* > 0 (the misalignment vector components *σ*′_*x*_ and *σ*′_*y*_), and 4 if *u* > 0 and *v* > 0 (the misalignment vector components *σ*_*x*_, *σ*_*y*_, *σ*′_*x*_ and *σ*′_*y*_), where the pre-factor *α*_*pqsuv*_ can be absorbed into the misalignment vectors in the cases where either *u* > 0 or *v* > 0 or both. Due to symmetry relations the actual number of independent parameters is actually less. Supplementary Table 2 summarizes all appearing terms up to and including order *j* = 2.

To lowest order *j* = 0 the aberration function only consists of defocus:

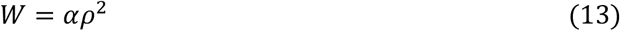

To first order *j* = 1 we find an aberration function that is a linear combination of the 8 additional invariants listed in Supplementary Table 2. We recognize one term related to primary spherical aberration, two terms related to primary coma, and the remaining 5 terms related to primary astigmatism and defocus/field curvature. The whole set of terms can be decomposed into Zernike modes:

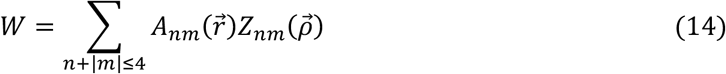

Where we list, for the sake of completeness, the appearing Zernike modes:

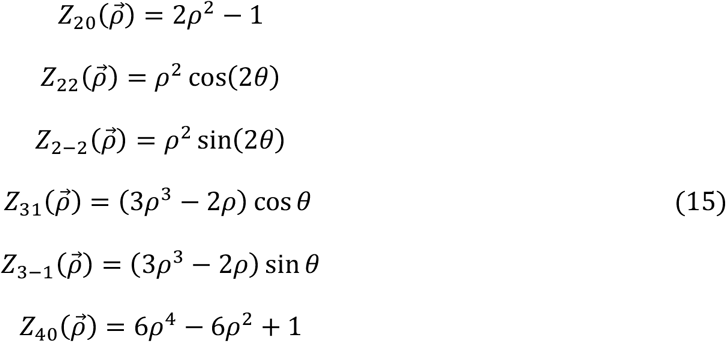

The participating terms per radial order *n* can be rewritten as follows. For primary spherical aberration we find (by adding an appropriate constant term and term in *ρ*^2^):

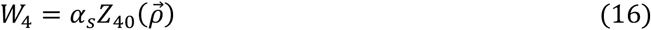

so that:

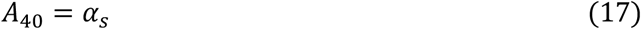

For primary coma we find:

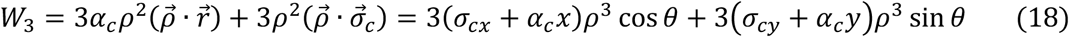

Adding tip/tilt (that can be added/subtracted at will since we will ignore this in the end anyway) gives Zernike coma coefficients:

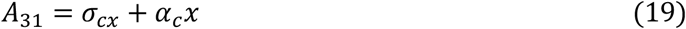

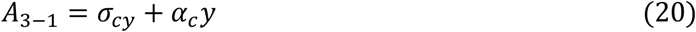

For primary astigmatism we can rearrange terms to obtain:

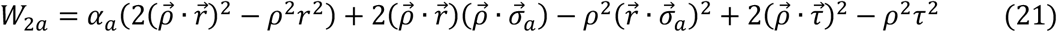

Where 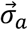 and 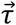 are misalignment vectors. The first term may be rewritten as:

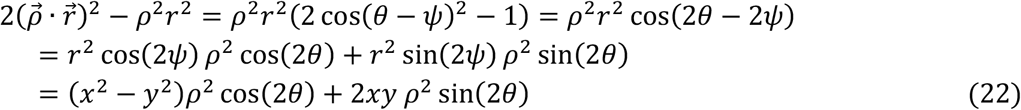

Similarly:

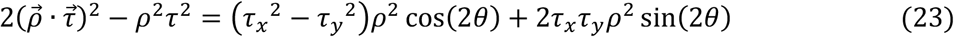

and:

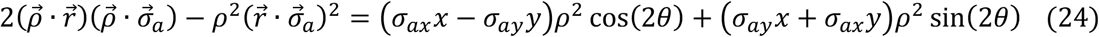

With *β*_22_ = *τ*_*x*_^2^ − *τ*_*y*_^2^ and *β*_2−2_ = 2*τ*_*x*_*τ*_*y*_ this gives Zernike astigmatism coefficients:

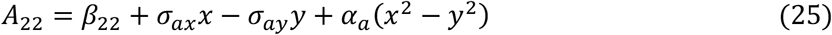

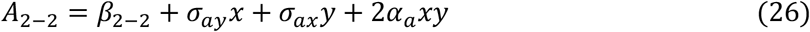

For defocus/field curvature, finally, we find:

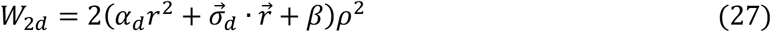

Adding piston we find that:

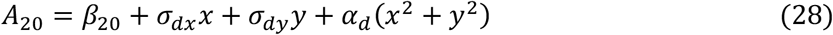

We can now count the number of parameters that describe the dependency of aberrations across the FOV. There are 4 parameters for defocus/field curvature 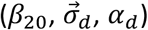, 5 parameters for astigmatism 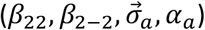, 3 parameters for coma 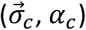, and 1 for spherical aberration (*α*_*s*_), giving a total of 13 parameters.

As all the aberration coefficients depend on the field coordinates up to 2nd order we can decompose them in terms of the Legendre polynomial products *Q*_00_(*x, y*), *Q*_10_(*x, y*), *Q*_01_(*x, y*), *Q*_20_(*x, y*), *Q*_11_(*x, y*), *Q*_02_(*x, y*). This finally gives the Legendre NAT coefficients as:

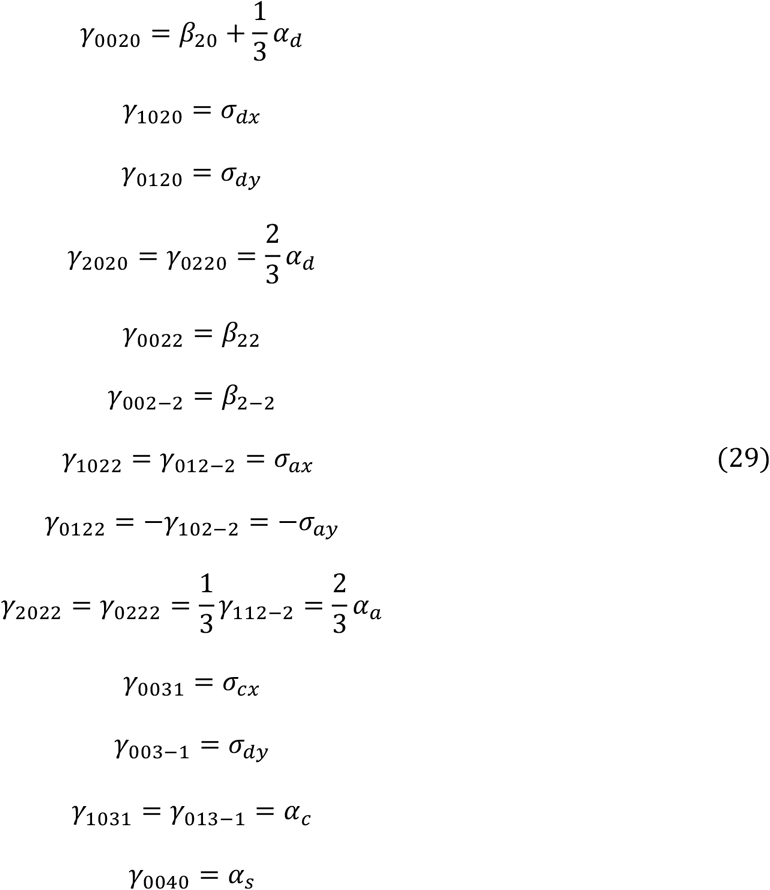

We see that there is a total of 19 non-zero Legendre NAT coefficients, that satisfy 6 symmetry relations, implying that there are 13 free parameters.

In order to link the 13 fit parameters to the system RMS aberration value we use the parametrization 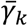 for *k* = 1,2 …, *N*_γ_ with *N*_γ_ = 13, and:

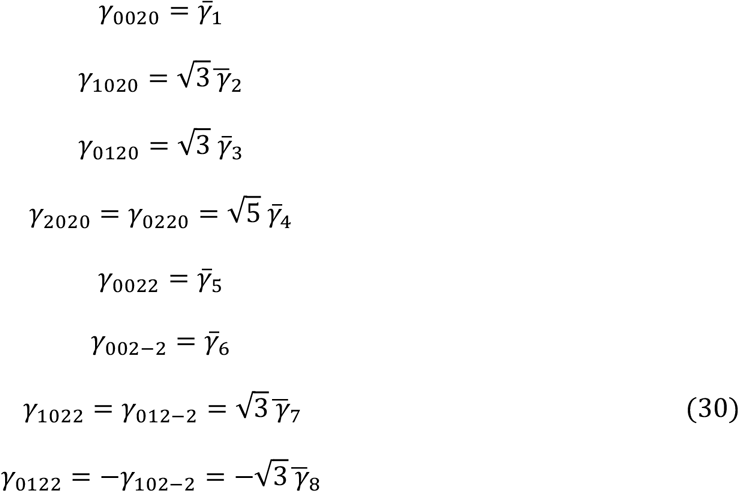

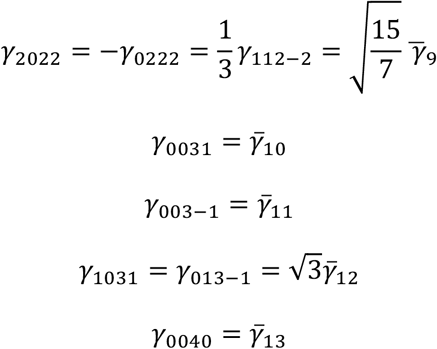

With this parametrization the system RMS aberration is given by:

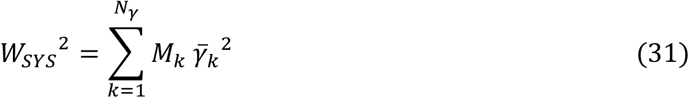

Here, *M*_*k*_ gives the multiplicity factor of each independent 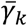, i.e. how many different Legendre NAT terms are proportional to 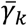. In this case *M*_9_ = 3, *M*_4_ = *M*_7_ = *M*_8_ = *M*_12_ = 2, and all other *M*_*k*_ = 1.

#### NAT for 2nd order aberration theory

In this case we have quite a bit more terms to cover, as there is an additional number of 25 invariants that have to be taken into account (see Supplementary Table 2). These invariants give rise to a total of 48 additional parameters, giving a total of 13+48 = 61 parameters. A careful analysis reveals that there is some redundancy in the invariants, reducing the number of free parameters to 56. The additional Zernike modes that appear up to this order are:

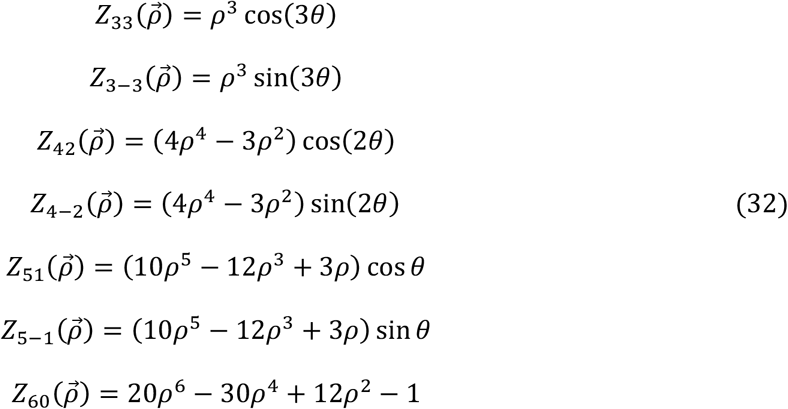

The field coordinates are expanded up to 4th order, implying we need the Legendre polynomial products *Q*_30_(*x, y*), *Q*_21_(*x, y*), *Q*_12_(*x, y*), *Q*_03_(*x, y*), *Q*_40_(*x, y*), *Q*_31_(*x, y*), *Q*_22_(*x, y*), *Q*_13_(*x, y*), and *Q*_04_(*x, y*) in addition to the first order aberration expansion. The appearing gamma NAT-coefficients and their symmetry relations can be derived similarly as for the first order expansion. The derivation of these is cumbersome and has been done with the help of Mathematica (See funextra/gammaNATcoefs.nb in Vectorfit [Vectorfit]). First the set of 9 existing invariants up to order *j* = 1 is expanded with 25 new relevant invariants at order *j* = 2 to a total of 34 invariants. Next, this set is divided into the order *n* of the pupil coordinates (see Supplementary Table 3). This gives 15 invariants with 29 parameters for *n* = 2 (defocus and primary astigmatism), 10 invariants with 19 parameters for *n* = 3 (primary coma and trefoil), 6 invariants with 9 parameters for *n* = 4 (primary spherical aberration and secondary astigmatism), 2 invariants with 3 parameters for *n* = 5 (secondary coma), and 1 invariant with 1 parameter for *n* = 6 (secondary spherical aberration). For each of these groups the invariants of Supplementary Table 2 are expanded in Legendre polynomial/Zernike mode products to find the set of non-zero Legendre NAT coefficients and to deduce the symmetry relations between these coefficients. This analysis shows that the number of truly free parameters is less than 61. The coma associated first order invariant 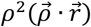 can be absorbed into the second order invariant 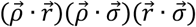, reducing the number of free parameters for the third order aberrations from 19 to 18. The defocus and astigmatism associated first order invariants 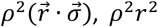, and 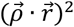 can likewise be absorbed into second order invariants, reducing the number of free parameters for the second order aberrations from 29 to 25. The final list of non-zero gamma NAT coefficients, grouped per aberration type, and expressed in the remaining total of *N*_γ_ = 56 free parameters 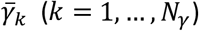 are listed in the following.

Second order terms (defocus, primary astigmatism): there are 15 invariants with 25 free parameters that give rise to 15 Legendre product polynomials for defocus and each type of primary astigmatism, so 45 terms in total, 8 of these are zero, and there are 12 symmetry relations, leading to 25 independent terms remaining:

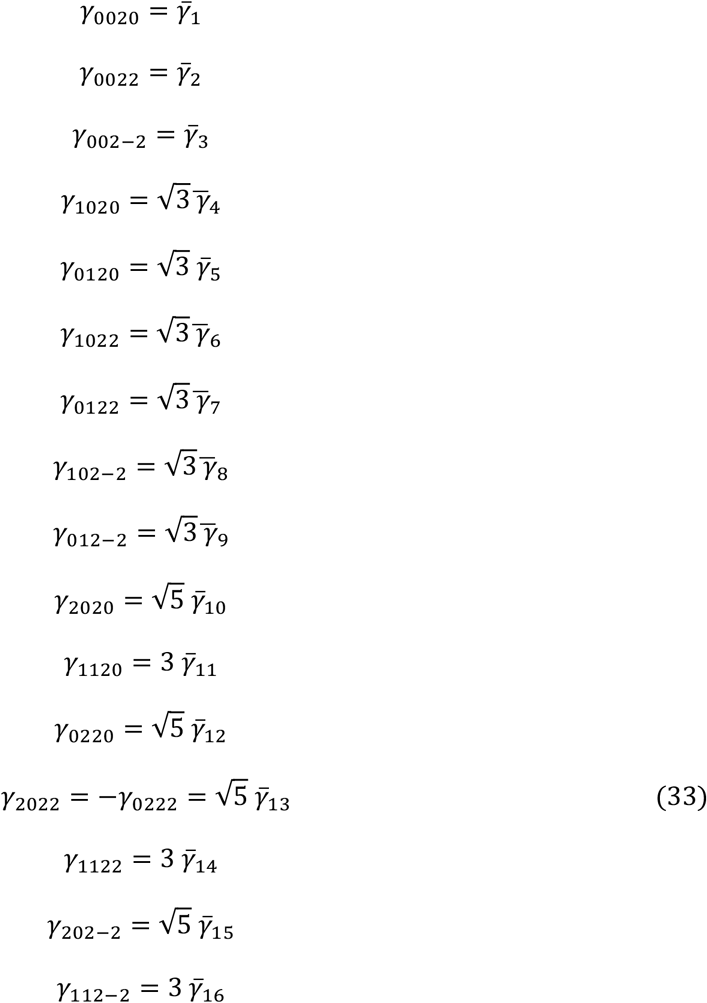

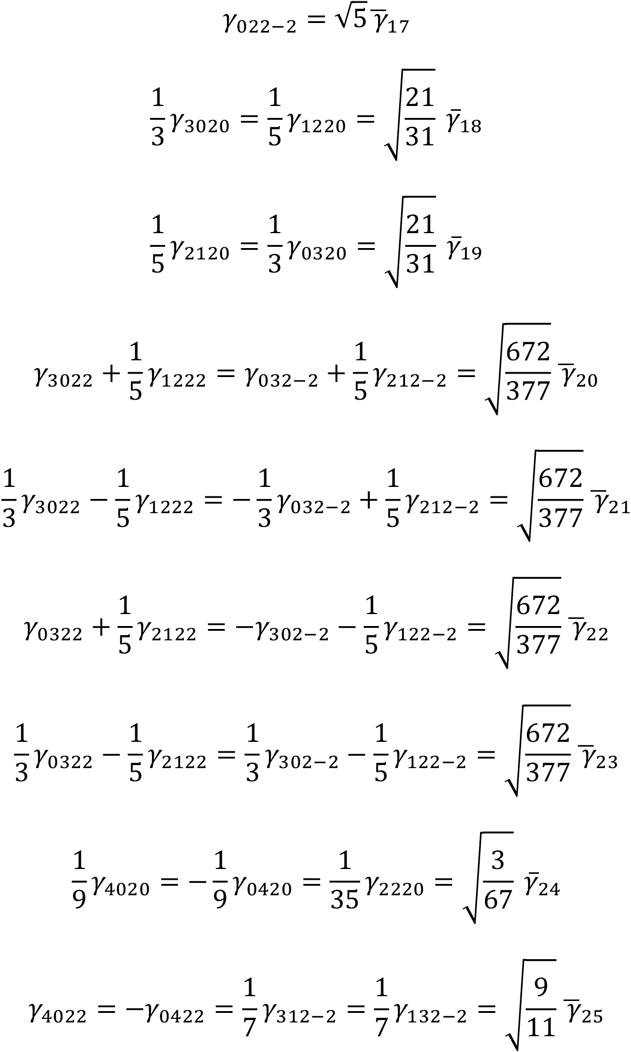

Symmetry relations with combined NAT coefficients can be inverted to:

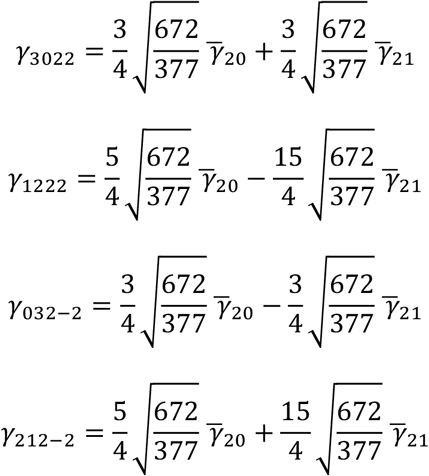

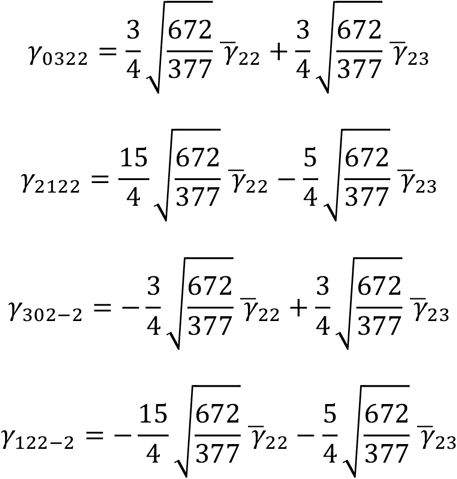

Third order terms (primary coma and trefoil): there are 10 invariants with 18 free parameters that give rise to 10 Legendre product polynomials for each type of primary coma and trefoil, so 40 terms in total, 8 of these are zero, and there are 14 symmetry relations, leading to 18 independent terms remaining. The 11 independent coefficients related to primary coma are:

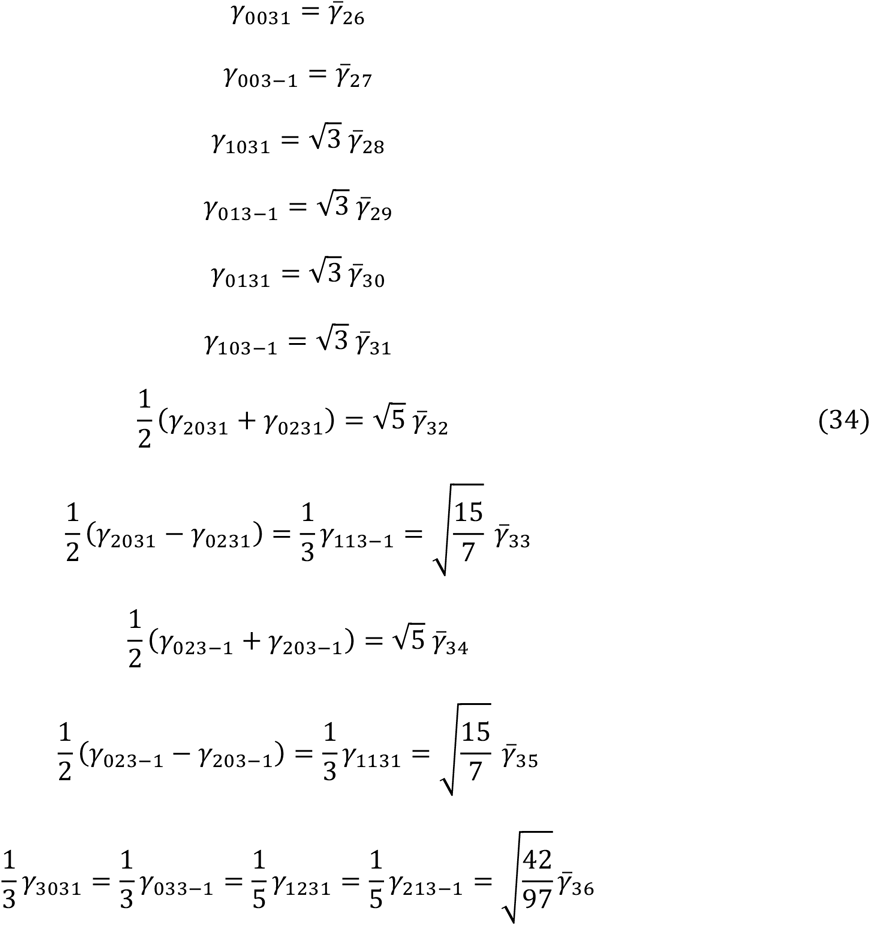

Symmetry relations with combined NAT coefficients can be inverted to:

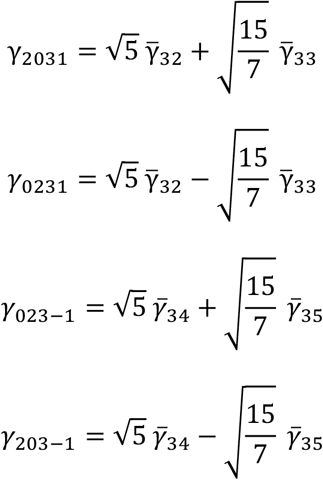

The 7 independent coefficients related to trefoil are:

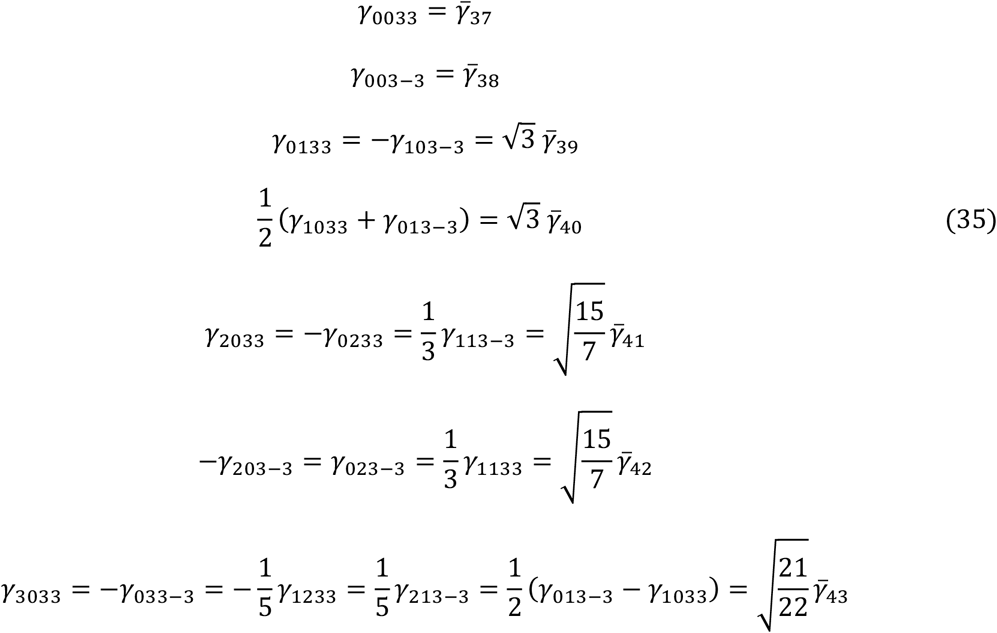

Symmetry relations with combined NAT coefficients can be inverted to:

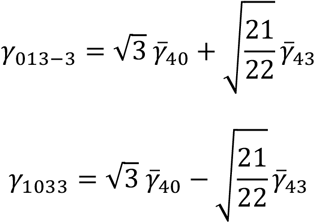

Fourth order terms (primary spherical aberration, secondary astigmatism): there are 6 invariants with 9 free parameters that give rise to 6 Legendre product polynomials for primary spherical aberration and each type of secondary astigmatism, so 18 different terms in total. Of these 4 are zero, there are 5 symmetry relations, leading to 9 independent terms remaining:

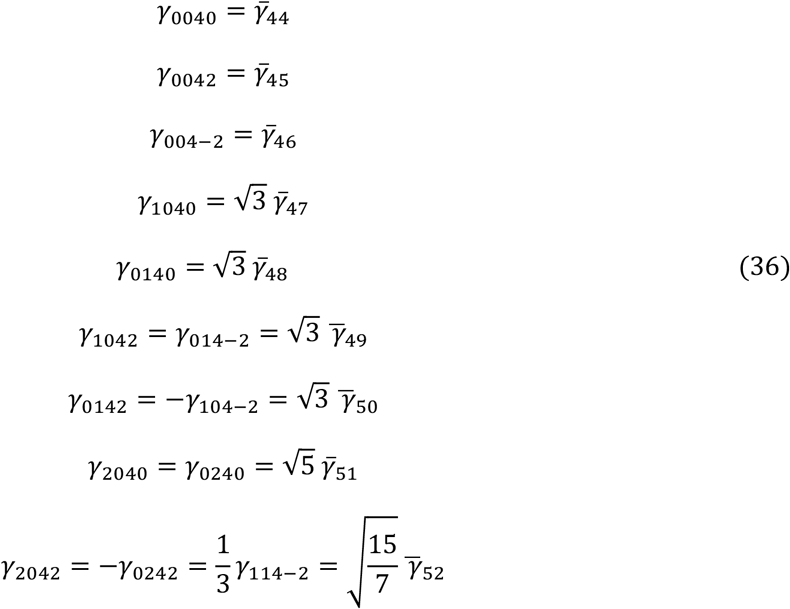

Fifth order terms (secondary coma): there are 2 invariants with 3 free parameters that give rise to 3 Legendre product polynomials for each type of secondary coma, so 6 different terms in total. Of these 2 are zero, there is 1 symmetry relation, leading to 3 independent terms remaining, with coefficients:

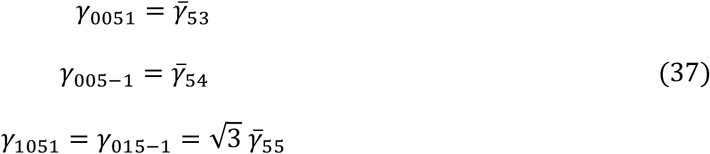

Sixth order term (secondary spherical aberration): there is only 1 invariant and 1 free parameter, that gives rise to only 1 Legendre product polynomial with coefficient:

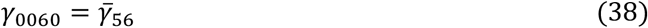

With this parameterization Eq. (31) still holds, where the new multiplicity factors are *M*_*k*_ = 2 for *k* = 13,18,19,32,34,39,40,49,50,51,55, *M*_*k*_ = 3 for *k* = 24,33,35,41,42,52, *M*_*k*_ = 4 for *k* = 20,21,22,23,25,36, *M*_*k*_ = 6 for *k* = 43, and *M*_*k*_ = 1 for all other *k*.

An alternative to the current approach to find the redundant set of NAT coefficients for higher orders is to arrange all participating gamma NAT coefficients in a vector *x* of length *N*, and rewrite the set of *K* symmetry relations as:

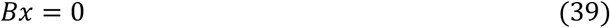

where *B* is a *K* × *N* matrix. The projection operator on the non-null space is:

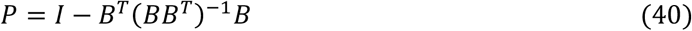

as *BP* = *B* − *B* = 0. Diagonalization of *P* leads to a set of eigenvectors *y*^(*l*)^, *l* = 1,2, …, *N* − *K* with eigenvalue 1, and in the optimization we could use an expansion of the gamma NAT coefficient vector *x* in terms of these eigenvector:

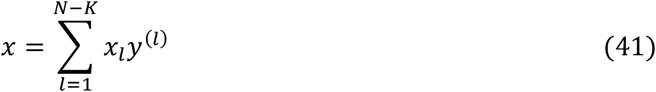

where the *x*_*l*_ can subsequently be used as independent optimization parameters.

**Supplementary Table 1:** Fitting speed of Vectorial PSF. Fits in 10^3^ fits/s for localizing a total of 1 million emitters that were simulated with the vectorial PSF model as described in the method section. The ROI size for 2D fitting is 7 × 7 and for 3D fitting 17 × 17. The GPU computations on the Nvidia A100 were performed using the DelftBlue supercomputer [DHPC2024]. The reported times include the computation of one OTF, assuming aberrations are constant over the FOV. When a grid of 20×20 OTFs is used, an additional 75 seconds is required for the computation.

|  | CPU using 1 core<br>Intel Xeon w5-2455X |  | CPU using all 12 cores<br>Intel Xeon w5-2455X |  | Desktop GPU<br>Nvidia RTX A4000 |  | High end HPC GPU<br>Nvidia A100 |  |
| --- | --- | --- | --- | --- | --- | --- | --- | --- |
|  | Direct | OTF | Direct | OTF | Direct | OTF | Direct | OTF |
| 2D | 0.6 | 1.5 | 4.0 | 8.7 | 17.4 | 61.5 | 50.3 | 100.0 |
| 3D | 0.2 | 0.5 | 1.4 | 2.7 | 6.3 | 6.8 | 14.8 | 10.1 |

**Supplementary Table 2:** NAT invariants and number of parameters. The table shows a list of all possible invariants of the form 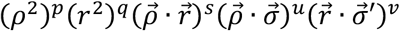 as defined in Eq. (10) for NAT order *j* = *p* + *q* + *s* + *u* + *v* − 1. The terms with overall order in the pupil coordinate 2*p* + *s* + *u* equal to 0 or 1 correspond to piston or wavefront tip/tilt and are ignored. These terms are marked with an asterisk.

| $j$ | $p$ | $q$ | $s$ | $u$ | $v$ | *invariant* | $2p+s+u$ | $2q+s+v$ | *#pars* |
| --- | --- | --- | --- | --- | --- | --- | --- | --- | --- |
| 0 | 1 | 0 | 0 | 0 | 0 | $\rho^2$ | 2 | 0 | 1 |
| | 0 | 1 | 0 | 0 | 0 | $r^2$ | 0* | 2 | 1 |
| | 0 | 0 | 1 | 0 | 0 | $\vec{\rho} \cdot \vec{r}$ | 1* | 1 | 1 |
| | 0 | 0 | 0 | 1 | 0 | $\vec{\rho} \cdot \vec{\sigma}$ | 1* | 0 | 2 |
| | 0 | 0 | 0 | 0 | 1 | $\vec{r} \cdot \vec{\sigma}$ | 0* | 1 | 2 |
| 1 | 2 | 0 | 0 | 0 | 0 | $\rho^4$ | 4 | 0 | 1 |
| | 0 | 2 | 0 | 0 | 0 | $r^4$ | 0* | 4 | 1 |
| | 0 | 0 | 2 | 0 | 0 | $(\vec{\rho} \cdot \vec{r})^2$ | 2 | 2 | 1 |
| | 0 | 0 | 0 | 2 | 0 | $(\vec{\rho} \cdot \vec{\sigma})^2$ | 2 | 0 | 2 |
| | 0 | 0 | 0 | 0 | 2 | $(\vec{r} \cdot \vec{\sigma})^2$ | 0* | 2 | 2 |
| | 1 | 1 | 0 | 0 | 0 | $\rho^2 r^2$ | 2 | 2 | 1 |
| | 1 | 0 | 1 | 0 | 0 | $\rho^2 (\vec{\rho} \cdot \vec{r})$ | 3 | 1 | 1 |
| | 1 | 0 | 0 | 1 | 0 | $\rho^2 (\vec{\rho} \cdot \vec{\sigma})$ | 3 | 0 | 2 |
| | 1 | 0 | 0 | 0 | 1 | $\rho^2 (\vec{r} \cdot \vec{\sigma})$ | 2 | 1 | 2 |
| | 0 | 1 | 1 | 0 | 0 | $r^2 (\vec{\rho} \cdot \vec{r})$ | 1* | 3 | 1 |
| | 0 | 1 | 0 | 1 | 0 | $r^2 (\vec{\rho} \cdot \vec{\sigma})$ | 1* | 2 | 2 |
| | 0 | 1 | 0 | 0 | 1 | $r^2 (\vec{r} \cdot \vec{\sigma})$ | 0* | 3 | 2 |
| | 0 | 0 | 1 | 1 | 0 | $(\vec{\rho} \cdot \vec{r})(\vec{\rho} \cdot \vec{\sigma})$ | 2 | 1 | 2 |
| | 0 | 0 | 1 | 0 | 1 | $(\vec{\rho} \cdot \vec{r})(\vec{r} \cdot \vec{\sigma})$ | 1* | 2 | 2 |
| | 0 | 0 | 0 | 1 | 1 | $(\vec{\rho} \cdot \vec{\sigma})(\vec{r} \cdot \vec{\sigma})$ | 1* | 1 | 4 |
| 2 | 3 | 0 | 0 | 0 | 0 | $\rho^6$ | 6 | 0 | 1 |
| | 0 | 3 | 0 | 0 | 0 | $r^6$ | 0* | 6 | 1 |
| | 0 | 0 | 3 | 0 | 0 | $(\vec{\rho} \cdot \vec{r})^3$ | 3 | 3 | 1 |
| | 0 | 0 | 0 | 3 | 0 | $(\vec{\rho} \cdot \vec{\sigma})^3$ | 3 | 0 | 2 |
| | 0 | 0 | 0 | 0 | 3 | $(\vec{r} \cdot \vec{\sigma})^3$ | 0* | 3 | 2 |
| | 2 | 1 | 0 | 0 | 0 | $\rho^4 r^2$ | 4 | 2 | 1 |
| | 2 | 0 | 1 | 0 | 0 | $\rho^4 (\vec{\rho} \cdot \vec{r})$ | 5 | 1 | 1 |
| | 2 | 0 | 0 | 1 | 0 | $\rho^4 (\vec{\rho} \cdot \vec{\sigma})$ | 5 | 0 | 2 |
| | 2 | 0 | 0 | 0 | 1 | $\rho^4 (\vec{r} \cdot \vec{\sigma})$ | 4 | 1 | 2 |
| | 1 | 2 | 0 | 0 | 0 | $\rho^2 r^4$ | 2 | 4 | 1 |
| | 0 | 2 | 1 | 0 | 0 | $r^4 (\vec{\rho} \cdot \vec{r})$ | 1* | 5 | 1 |
| | 0 | 2 | 0 | 1 | 0 | $r^4 (\vec{\rho} \cdot \vec{\sigma})$ | 1* | 4 | 2 |
| | 0 | 2 | 0 | 0 | 1 | $r^4 (\vec{r} \cdot \vec{\sigma})$ | 0* | 5 | 2 |
| | 1 | 0 | 2 | 0 | 0 | $\rho^2 (\vec{\rho} \cdot \vec{r})^2$ | 4 | 2 | 1 |
| | 0 | 1 | 2 | 0 | 0 | $r^2 (\vec{\rho} \cdot \vec{r})^2$ | 2 | 4 | 1 |
| | 0 | 0 | 2 | 1 | 0 | $(\vec{\rho} \cdot \vec{r})^2 (\vec{\rho} \cdot \vec{\sigma})$ | 3 | 2 | 2 |
| | 0 | 0 | 2 | 0 | 1 | $(\vec{\rho} \cdot \vec{r})^2 (\vec{r} \cdot \vec{\sigma})$ | 2 | 3 | 2 |
| | 1 | 0 | 0 | 2 | 0 | $\rho^2 (\vec{\rho} \cdot \vec{\sigma})^2$ | 4 | 0 | 2 |
| | 0 | 1 | 0 | 2 | 0 | $r^2 (\vec{\rho} \cdot \vec{\sigma})^2$ | 2 | 2 | 2 |
| | 0 | 0 | 1 | 2 | 0 | $(\vec{\rho} \cdot \vec{r})(\vec{\rho} \cdot \vec{\sigma})^2$ | 3 | 1 | 2 |
| | 0 | 0 | 0 | 2 | 1 | $(\vec{\rho} \cdot \vec{\sigma})^2 (\vec{r} \cdot \vec{\sigma})$ | 2 | 1 | 4 |
| | 1 | 0 | 0 | 0 | 2 | $\rho^2 (\vec{r} \cdot \vec{\sigma})^2$ | 2 | 2 | 2 |
| | 0 | 1 | 0 | 0 | 2 | $r^2 (\vec{r} \cdot \vec{\sigma})^2$ | 0* | 3 | 2 |
| | 0 | 0 | 1 | 0 | 2 | $(\vec{\rho} \cdot \vec{r})(\vec{r} \cdot \vec{\sigma})^2$ | 1* | 3 | 2 |
| | 0 | 0 | 0 | 1 | 2 | $(\vec{\rho} \cdot \vec{\sigma})(\vec{r} \cdot \vec{\sigma})^2$ | 1* | 2 | 4 |
| | 1 | 1 | 1 | 0 | 0 | $\rho^2 r^2 (\vec{\rho} \cdot \vec{r})$ | 3 | 3 | 1 |
| | 1 | 1 | 0 | 1 | 0 | $\rho^2 r^2 (\vec{\rho} \cdot \vec{\sigma})$ | 3 | 2 | 2 |
| | 1 | 0 | 1 | 1 | 0 | $\rho^2 (\vec{\rho} \cdot \vec{r})(\vec{\rho} \cdot \vec{\sigma})$ | 4 | 1 | 2 |
| | 0 | 1 | 1 | 1 | 0 | $r^2 (\vec{\rho} \cdot \vec{r})(\vec{\rho} \cdot \vec{\sigma})$ | 2 | 3 | 2 |
| | 1 | 1 | 0 | 0 | 1 | $\rho^2 r^2 (\vec{r} \cdot \vec{\sigma})$ | 2 | 3 | 2 |
| | 1 | 0 | 1 | 0 | 1 | $\rho^2 (\vec{\rho} \cdot \vec{r})(\vec{r} \cdot \vec{\sigma})$ | 3 | 2 | 2 |
| | 0 | 1 | 1 | 0 | 1 | $r^2 (\vec{\rho} \cdot \vec{r})(\vec{r} \cdot \vec{\sigma})$ | 1* | 4 | 2 |
| | 1 | 0 | 0 | 1 | 1 | $\rho^2 (\vec{\rho} \cdot \vec{\sigma})(\vec{r} \cdot \vec{\sigma})$ | 3 | 1 | 4 |
| | 0 | 1 | 0 | 1 | 1 | $r^2 (\vec{\rho} \cdot \vec{\sigma})(\vec{r} \cdot \vec{\sigma})$ | 1* | 3 | 4 |
| | 0 | 0 | 1 | 1 | 1 | $(\vec{\rho} \cdot \vec{r})(\vec{\rho} \cdot \vec{\sigma})(\vec{r} \cdot \vec{\sigma})$ | 2 | 2 | 4 |

**Supplementary Table 3:** Number of free parameters per radial aberration order. The table shows the number of invariants and associated number of parameters per radial aberration order *n* and per NAT order *j* as deduced from Supplementary Table 2. Due to redundancy in the lower order invariants when going to order *j* = 2, the number of free parameters is reduced.

| $n$ | # invariants | | | | # parameters | | | | |
| --- | --- | --- | --- | --- | --- | --- | --- | --- | --- |
| | $j=0$ | $j=1$ | $J=2$ | *total* | $J=0$ | $J=1$ | $J=2$ | *total* | *independent* |
| 2 | 1 | 5 | 9 | 15 | 1 | 8 | 20 | 29 | 25 |
| 3 | 0 | 2 | 8 | 10 | 0 | 3 | 16 | 19 | 18 |
| 4 | 0 | 1 | 5 | 6 | 0 | 1 | 8 | 9 | 9 |
| 5 | 0 | 0 | 2 | 2 | 0 | 0 | 3 | 3 | 3 |
| 6 | 0 | 0 | 1 | 1 | 0 | 0 | 1 | 1 | 1 |
| *total* | 1 | 8 | 25 | 34 | 1 | 12 | 48 | 61 | 56 |

## Notes

### Competing Interest Statement

The authors have declared no competing interest.

